# Peptide additives reprogram the lipid nanoparticle corona and enhance gene delivery in a serum-free environment for lung epithelium

**DOI:** 10.64898/2026.07.10.737795

**Authors:** Jinzhen Hu, Michael B. Papah, Arlett Ramirez, Deepthi Alapati, Millicent O. Sullivan

## Abstract

Lipid nanoparticles (LNPs) have become a clinical standard for systemically-administered nucleic acid drugs and vaccines, but the LNP pipeline for locally-delivered LNP therapies remains much less mature. Local delivery in lung represents a particularly compelling application space for DNA-LNP therapeutics, as local gene therapies could support sustained epithelial recovery and functional restoration in various lung diseases. However, locally-delivered DNA-LNPs face multiple barriers, including the limited availability of serum components that often support conventional LNP activity, and the additional delivery barriers posed by the nucleus. We generated hybrid peptide-lipid nanoparticles (hpLNPs) for pulmonary DNA delivery by using a core-shell assembly strategy to incorporate short histone-derived peptides, selected for their DNA-transport capacity, into a clinically inspired LNP formulation. In parallel, we evaluated serum pre-coating of LNPs as a strategy to boost hpLNP activity in the serum-poor airway environment. A peptide:DNA amine/phosphate (N/P) ratio of 0.9 was identified as the highest feasible ratio to permit peptide incorporation into hpLNPs while preserving DNA encapsulation efficiency at >90%, retaining hpLNP colloidal stability, and preserving the overall pKa for LNPs. At N/P = 0.9, hpLNPs showed markedly enhanced DNA delivery in alveolar lung cells, achieving up to a 17-fold increase in transgene expression compared to peptide-free LNPs. Transgene expression levels varied depending on serum concentration, with expression peaking in the presence of 6% serum. Furthermore, serum pre-coating was necessary to enable *in vivo* hpLNP activity following intratracheal administration in mice, yielding robust local GFP expression. Mechanistic studies in exosome-free serum revealed that peptide incorporation in the hpLNPs enhanced transgene delivery by increasing nanoparticle interactions with serum exosome components, resulting in significant enhancements to hpLNP uptake. Together, these findings identify nanoparticle– serum interactions as a critical determinant of LNP-mediated pulmonary DNA delivery and establish peptide incorporation and serum pre-coating as promising strategies to enable robust, localized gene transfer in serum-free environments.

## Introduction

Lipid nanoparticles (LNPs) have become the leading platform for nucleic acid delivery, building on the successful launch of the mRNA-based COVID-19 vaccines.^1,2^ More broadly, the LNP therapeutic pipeline has expanded substantially, spanning preclinical candidates, clinical-stage programs, and FDA-approved medicines, with most progress to date centered on mRNA and siRNA cargos for cancer, neurodegenerative diseases, and rare genetic or metabolic disorders.^3–11^ In contrast, DNA delivery via LNPs remains comparatively less developed, though DNA offers distinct advantages for some applications. For example, like mRNA, DNA-based approaches can directly restore expressions of missing or defective proteins, thereby addressing the underlying dysfunction. In addition, DNA therapeutics can produce more prolonged expression than inherently less stable mRNA drugs, offering pathways to reduce dosing frequency and provide valuable alternative treatment strategies in chronic diseases where durable correction is needed.^12^ DNA vectors also can accommodate larger and more complex design features, and the inherent chemical properties of DNA can induce immunostimulatory effects relevant in the context of vaccination, expanding the therapeutic scope.^13,14^ From a manufacturing and distribution perspective, similar to current mRNA based products, DNA platforms support rapid, scalable manufacturing, and DNA vectors can additionally offer practical advantages in storage and distribution because of the greater physicochemical stability of DNA and reduced reliance on ultra-cold-chain logistics.^15–18^

Building on the success of LNPs as delivery vehicles for mRNA and siRNA, researchers have begun to address key barriers limiting DNA delivery. In the cargo engineering context, plasmid constructs have been engineered with scaffold/matrix attachment regions (S/MARs) to permit sustained nuclear retention and improve expression durability, enabling prolonged generation of recombinant T cells.^19,20^ In the formulation context, endogenous anti-inflammatory lipids that inhibit the DNA stimulator of interferon genes (STING) pathway have been incorporated into LNPs to reduce intracellular inflammatory signaling triggered by foreign DNA, particularly for non-vaccine applications.^21^ Despite these advances, studies on non-lipid additives—which could introduce new functionalities, address the challenge of nuclear uptake, and refine design principles related to encapsulation and charge interactions—remain scarce in the DNA–LNP space.

The lung represents a compelling application space for DNA–LNP therapeutics, particularly within the alveolar compartment, where homeostasis depends on coordinated interactions between alveolar epithelial cells and resident macrophages.^22^ Among the cell types that reside in the alveolar region, alveolar epithelial cells such as alveolar type 2 progenitor (AT2) cells are of particular interest because they produce surfactants, support alveolar immune homeostasis, and serve as the principal progenitor population responsible for epithelial repair through differentiation toward the alveolar type 1 (AT1) lineage.^23–28^ Disruption of AT2-mediated repair contributes to fibrotic lung diseases such as pulmonary fibrosis, in which defective epithelial regeneration and barrier restoration drive progressive remodeling.^29^ In this context, DNA delivery is attractive because its longer-lasting transgene expression may better support sustained epithelial recovery and a regain of function.^30,31^ These considerations also motivate local pulmonary administration, which can increase exposure at the site of pathology while limiting systemic off-target effects. However, LNP formulation performance in the alveoli cannot be inferred directly from intravenous performance, since LNPs in blood rapidly acquire biomolecular coronas that drive biodistribution and cellular uptake, whereas the alveolar environment has distinct chemical and biological components.^32,33^ As a result, locally-delivered LNPs encounter a distinct microenvironment that may not provide the same corona-dependent behavior exhibited by systemically-delivered LNP formulations. Although prior studies of inhaled gene delivery have focused extensively on extracellular barriers such as airway mucus, particularly in obstructive lung disease, an additional challenge for alveolar delivery is understanding how LNPs perform under serum-free conditions and whether favorable LNP–biomolecule interactions must be engineered or deliberately reintroduced for effective local delivery in lung.

Expanding our previous work showing that histone 3 (H3)-derived peptides (containing the amino acid sequence derived from the H3 tail, positions 1-25) can enhance nuclear uptake and DNA unpackaging in polymer nanostructures, we sought to test whether H3 peptides also offered these benefits within a clinically-relevant LNP platform,^34–36^ and whether peptide-modified LNPs could facilitate efficient gene delivery in the serum-poor lung environment. Using an LNP formulation containing the ionizable lipid ALC-0315, and employing a lipid composition derived from Pfizer mRNA coronavirus vaccine as a benchmark, we incorporated the H3 peptide using a core-shell assembly method to pre-complex DNA with H3 peptides and subsequently add a lipid shell.^37^ Serum free transfection studies in murine lung epithelial (MLE-12) cells showed that both regular LNPs (rLNPs) and hybrid peptide-lipid nanoparticles (hpLNPs) produced minimal transgene expression. Likewise, no transgene expression was detected following intratracheal delivery in mice. However, serum pre-coating enabled expression both *in vitro* and *in vivo* for both types of LNPs. In particular, the combinatorial effect in gene expression from serum pre-coating and peptide inclusion was substantial, achieving up to a 17-fold increase in gene expression relative to rLNPs *in vitro*. While we hypothesized that peptide incorporation would boost transgene expression by leveraging H3 effectors to enhance nuclear delivery, hpLNPs containing scrambled-H3 (sH3) peptides exhibited similar behavior as hpLNPs containing H3 peptides, demonstrating that the effect was sequence independent. Consistent with this observation, cell and nuclear uptake assessments with Cy5-stained hpLNPs revealed that peptide incorporation enhanced cellular uptake by 20–30-fold but did not increase nuclear accumulation. Further studies using exosome-depleted serum suggested that interactions with serum-derived exosome components were key drivers of cellular uptake and subsequent gene expression. These findings identify serum association as a critical determinant of LNP-mediated gene delivery in the serum-poor lung environment, and highlight serum precoating and peptide incorporation as effective strategies to advance efficacy in LNP platforms for pulmonary gene therapy applications.

## Results and discussion

### Core-shell assembly and characterization of hpLNPs

hpLNPs were designed and formulated using a core-shell assembly strategy to generate nanoparticles consisting of an inner peptide–DNA complex and an outer lipid shell (Figure 1). Because the histone-tail peptide is cationic, its incorporation was expected to alter the electrostatic interactions between the DNA complex and the lipid layer during microfluidic mixing, thereby influencing both the hpLNP particle size and the quantity and composition of the lipid coating. The range of peptide:DNA N/P ratios tested (with positive charges from peptide amines and negative charges from DNA phosphates) was selected to balance two considerations: maintaining a slightly anionic inner complex to preserve the electrostatic driving force for subsequent lipid encapsulation, and maximizing H3 loading levels. N/P ratios greater than 1 were additionally included to examine how departure from this assembly-favorable regime influenced nanoparticle formation. A clinically-derived lipid composition was selected as the benchmark system (stemming from formulations reported in the Pfizer COVID vaccine patent).^37^ Particle size and encapsulation efficiency were evaluated by dynamic light scattering (DLS) and gel electrophoresis, respectively.

**Figure 1.**
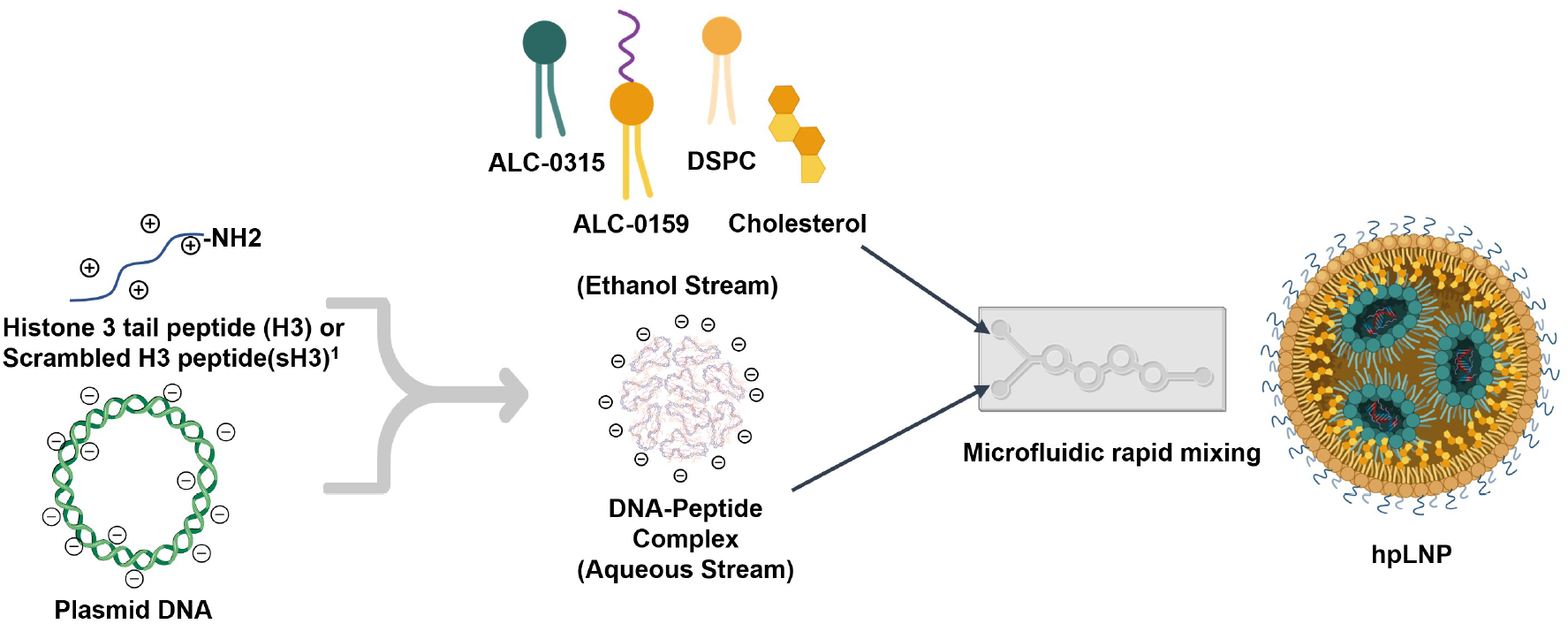
Schematic diagram of the stepwise formulation process to produce the peptide-DNA hpLNP.

The peptide:DNA precomplex was produced by vortex mixing. Following precomplex assembly, lipid components were added to the precomplexed core through microfluidic mixing. Gel electrophoresis on the resulting LNPs or hpLNPs demonstrated a complete disappearance of the DNA bands, consistent with ethidium bromide exclusion in intact nanoparticles (Figure 2A). Gel electrophoresis following treatment with detergent to disassemble the lipid coating (Figure 2A, right) showed a progressive reduction in DNA band migration with increasing peptide:DNA N/P ratio, indicating sustained peptide–DNA association throughout the core-shell assembly process. DNA-peptide binding was preserved up to a peptide:DNA N/P ratio of 0.9, but decreased markedly at ratios above 1. This behavior was further supported by DLS analysis: at lower N/P ratios, hpLNPs exhibited the expected size increase associated with successful encapsulation of the peptide–DNA core in addition to the LNP coating, whereas at higher N/P ratios the average particle size shifted back toward values comparable to empty LNPs, suggesting inefficient encapsulation of the pre-complexed cargo (Figure 2B). Furthermore, formulations prepared above N/P = 1 also displayed broader secondary DLS peaks, consistent with the formation of a separate and less stable peptide–DNA aggregate population rather than a single hybrid nanoparticle population. These results suggest that the decline in encapsulation as the core approached and exceeded charge neutrality likely reflected a reduced electrostatic driving force for controlled ionizable lipid encapsulation of the peptide–DNA complex. As this driving force weakened, the assembly process became less effective, resulting in enhanced aggregation and lowering overall encapsulation efficiency. A similar charge-dependent trend was observed when nanoparticles were formed through pipette-based core–shell assembly, but the loss in encapsulation was more pronounced as the core approached charge neutrality (Figure S1 and Table S1). This suggested that peptide–DNA core formulations near neutral charge are particularly sensitive to the less controlled and more heterogeneous mixing environment associated with manual pipette mixing. In contrast, the more defined and reproducible mixing conditions provided by microfluidic assembly appeared to better preserve the controlled lipid deposition onto the peptide–DNA core. Collectively, these results identified N/P = 0.9 as the highest peptide loading condition that still supported stable hpLNP assembly and efficient encapsulation, and accordingly, this formulation was selected for further assessment.

**Figure 2.**
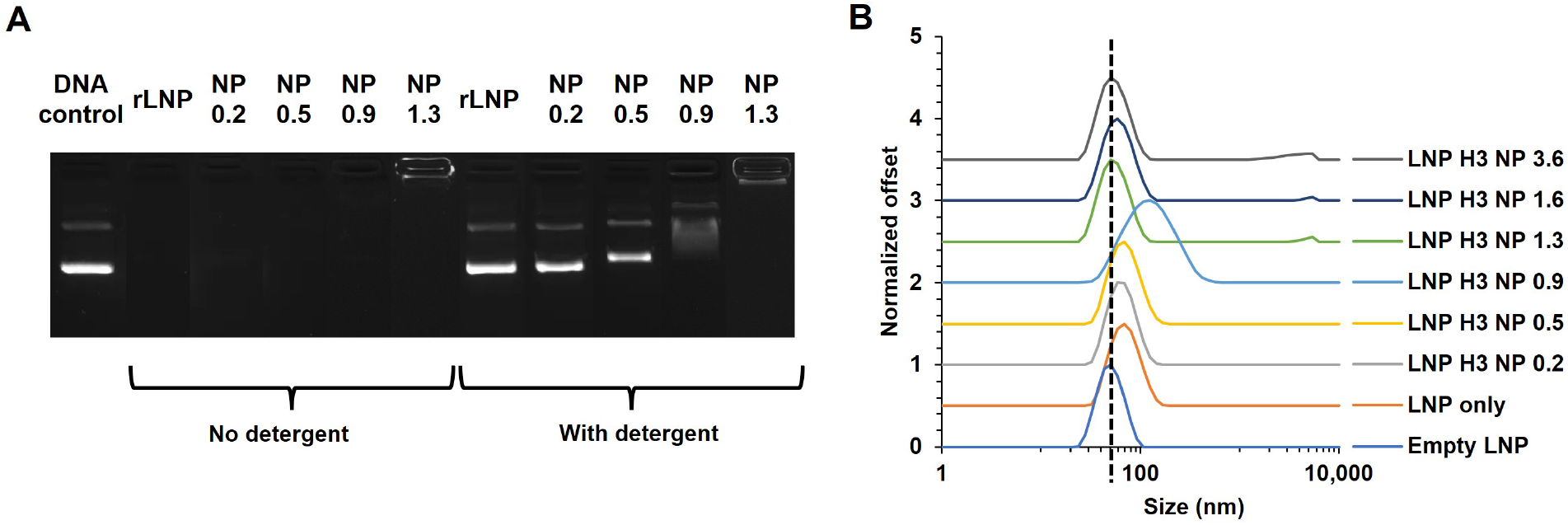
Assessment of peptide incorporation efficacy and stability. (A) Representative agarose gel assessment of peptide-DNA encapsulation and complexation of peptide-DNA precomplexes at different N/P ratios. (B) DLS size measurements on LNPs with different N/P ratios and quantities of peptide incorporated. The dashed line indicates the size of the empty LNPs.

To establish whether the selected N/P = 0.9 condition remained compatible with LNP formulation requirements, hpLNPs containing either H3 peptides or an identical quantity of sH3 peptides was subjected to systematic physicochemical characterization. Peptide incorporation led to modest increases in particle size and polydispersity index (PDI) relative to peptide-free controls (Figure 3A, B, and D), but all formulations remained within a size range generally considered suitable for LNP delivery (∼60–150 nm).^38^ PicoGreen measurements showed a slight reduction in DNA encapsulation efficiency upon peptide inclusion; however, all formulations retained greater than 90% encapsulation, indicating that peptide incorporation had minimal impact on nucleic acid loading capacity (Figure 3A–D). Likewise, the apparent pKa values of hpLNPs (6.60–6.65) were nearly identical to those of peptide-free controls, suggesting that the pH-responsive ionization behavior of the ionizable lipid was preserved despite peptide incorporation (Supplemental Figure S2 and Table S2). Triton X addition combined with heparin displacement restored DNA migration in hpLNPs, further confirming that peptide–DNA interactions were maintained during formulation and remained reversible under competitive polyanion challenge (Figure 3E).

**Figure 3.**
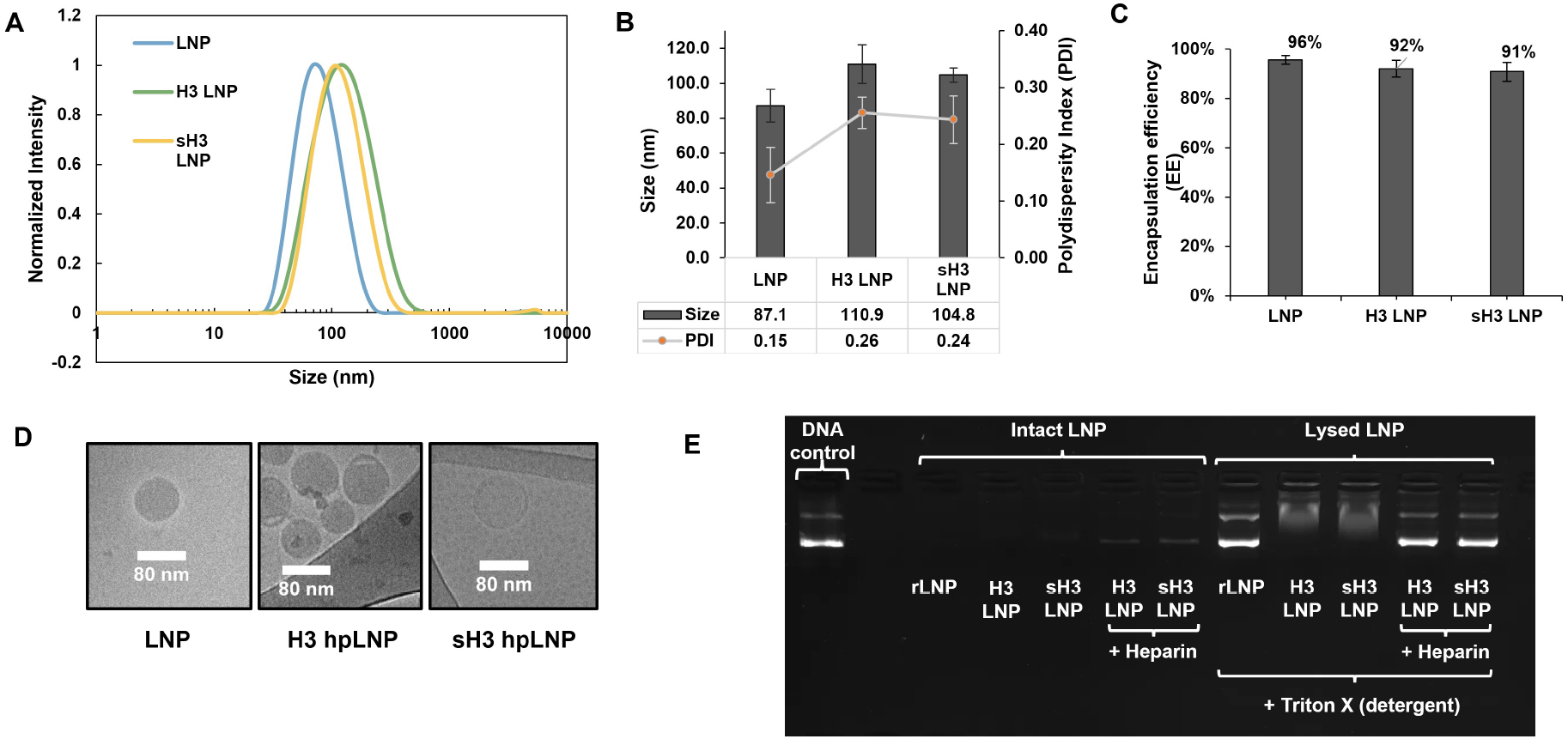
Physical characterization of the different kinds of LNPs. (A, B) DLS size measurements on all three types of LNPs, with all hpLNPs formulated at an N/P ratio of 0.9 (n=6). (C) Encapsulation efficiency measurements on all three types LNPs (n=6). (D) Representative images made using cryogenic electron microscopy (cryo-EM) on all three types of LNPs. (E) Representative agarose gel electrophoresis showing the degree of LNP encapsulation and cargo complexation via ethidium bromide staining.

### Serum-dependent transfection of LNPs in MLE-12 lung cells and murine lung epithelium

Because the protein corona is known to strongly influence LNP uptake and biodistribution, its absence may substantially alter LNP–cell interactions.^39^ However, serum proteins do not exert a consistent effect on LNP delivery; their impact depends on lipid composition, cell type, cargo, and biological context, with reported outcomes ranging from enhanced delivery to reduced activity.^40–43^ This context dependence highlights the need to evaluate serum effects directly, particularly when LNP composition is modified. We therefore examined the transfection activity of both rLNPs and hpLNPs *in vitro*, under serum-free conditions, and *in vivo* after local pulmonary administration in mice. Under serum-free culture, both formulations produced minimal GFP expression, as shown by fluorescence imaging and confirmed by flow cytometry (Figure 4A). Although two days of serum-free culture may slow proliferation and reduce plasmid DNA expression in cells, the cells remained viable and continued to divide, indicating that their capacity for transgene expression was not abolished in an obvious way. Furthermore, when rLNPs and hpLNPs were delivered via intratracheal administration in mouse lungs, a similar lack of detectable GFP expression was observed in the lung as compared to the levels of GFP expression in mouse genetically modified to express GFP (Figure 4B). The loss of expression in serum-free conditions (both *in vitro* and *in vivo*), most likely reflects impaired LNP-cell interactions rather than the effect of serum starvation on cell function alone. These findings suggest that serum proteins are critical for effective LNP uptake and subsequent gene expression, likely due to their role in forming a protein corona that facilitates cell interaction.

**Figure 4.**
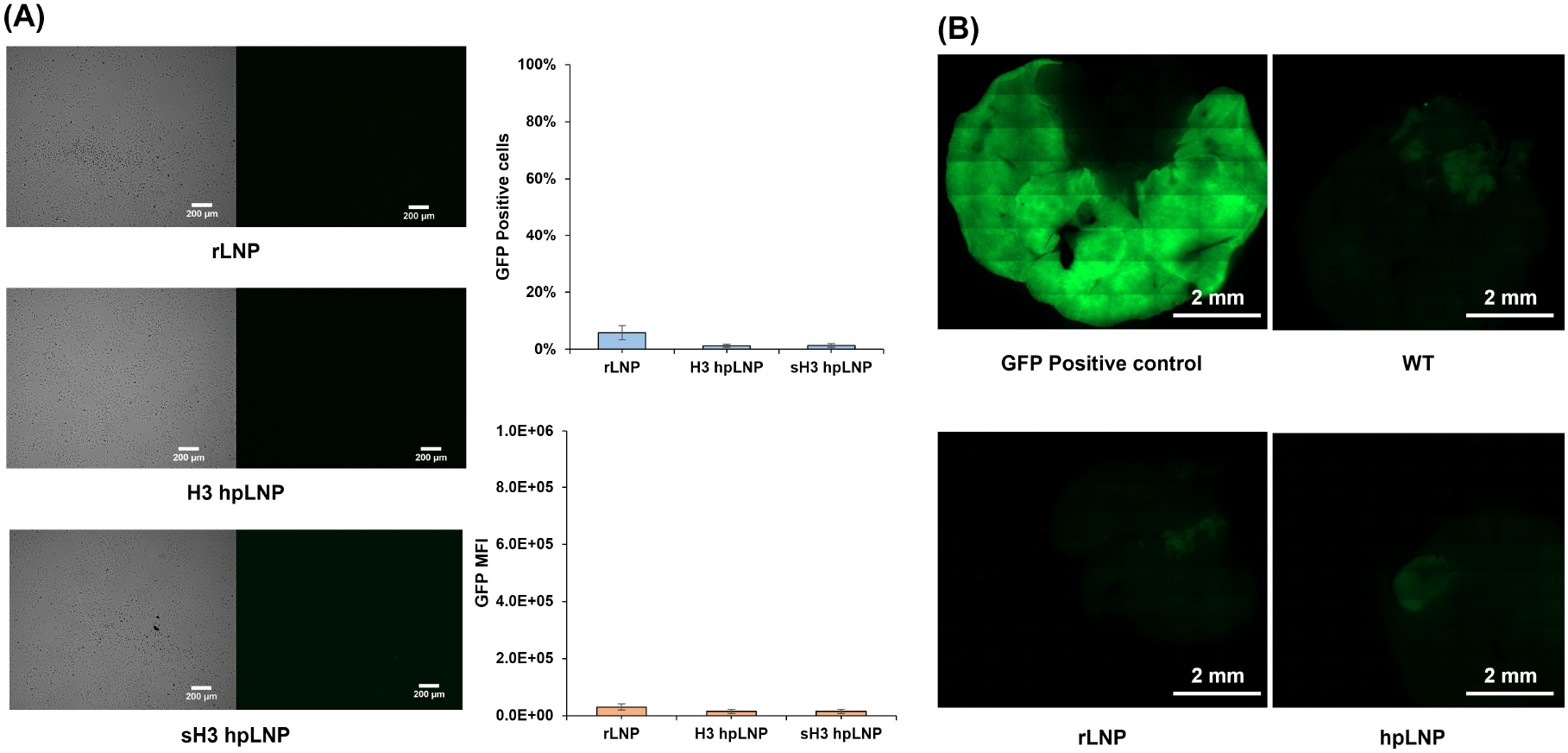
LNP transfection in serum free environments. (A) *In vitro* delivery of various LNPs in MLE-12 cells. Representative phase contrast (far left) and fluorescent (center left) images of MLE-12 cells 48 h after treatment with various LNP formulations under serum free conditions. Corresponding quantification of GFP expression in transfected MLE-12 cells based on flow cytometry analysis of transfection efficiency (right, top) and overall gene expression (right, bottom) (n=3). (B) Intratracheal delivery of rLNPs and hpLNPs (bottom) as compared with GFP positive control lungs and GFP negative control lungs from wild type (WT), untreated mouse (top). Representative tiled fluorescent images of murine lungs taken 2 days after LNP administration.

For polymeric cationic nanoparticles, poor serum stability is a limitation for prolonged dosing.^44,45^ To evaluate whether our hybrid LNPs faced a similar challenge, we first assessed their stability in serum-containing media prior to uptake and transfection experiments. All formulations were incubated in growth media containing 2–10% fetal bovine serum (FBS), and particle size was monitored over time by DLS at 0, 7, 16, 30, and 44 h (Figure 5 and Supplemental Figure S3). All three LNP types exhibited an initial size increase upon serum exposure, and LNP size continued to increase gradually until approximately 16 hours, after which it plateaued and remained stable through 44 hours. Importantly, no evidence of aggregation was observed, indicating that both rLNPs and hpLNPs maintained colloidal stability during prolonged serum exposure. These results are consistent with an initial rapid serum component association on the LNP surface, followed by a saturation of serum components on the LNP surface after approximately 16 hours. To provide a consistent platform for assessing serum effects on cellular transfection, all assessments of LNPs in serum-containing conditions were performed following LNP pre-incubation in serum for 24 h prior to cell dosing to ensure that the particles had reached a stable, serum-conditioned state.

**Figure 5.**
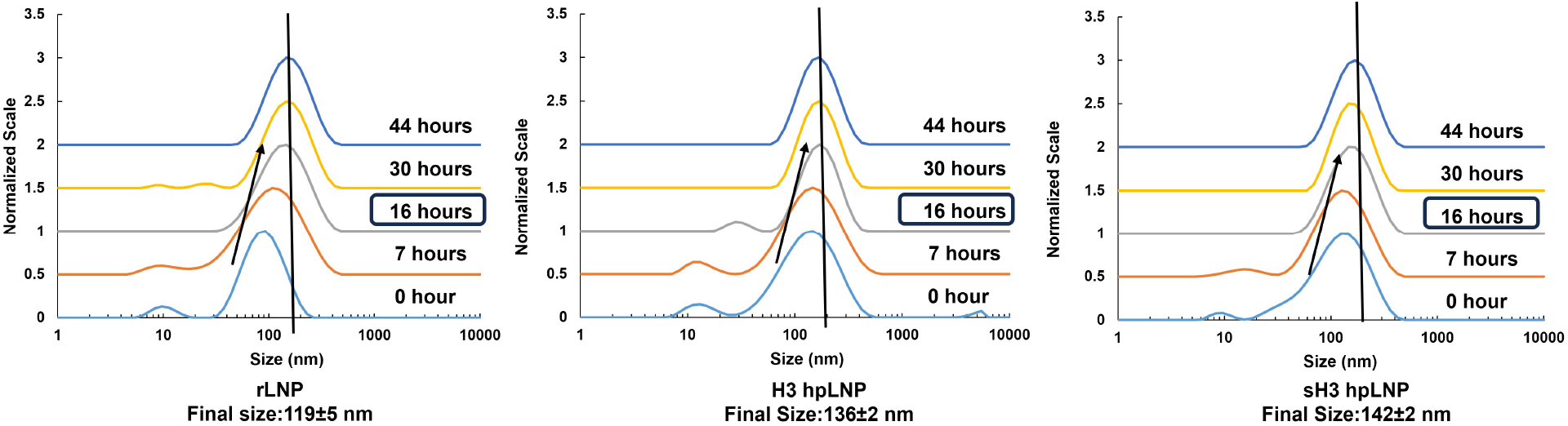
Representative time course DLS size measurements of various LNPs in full serum media. The initial (t=0) size measurements were taken immediately after LNP addition into the media. The arrow shows the progression in size of the LNPss and the line indicates the time point at which no further evolution in size occurred.

Having established that prolonged serum exposure altered LNP hydrodynamic size without inducing aggregation, we next asked whether the serum-conditioned particles induced efficient gene delivery. rLNPs, H3-hpLNPs, and sH3 hpLNPs were pre-incubated with growth media containing 2–10% FBS for 24 h at 37 °C and then the particles were dosed onto MLE-12 cells for 48 h. GFP expression was quantified by fluorescence imaging and flow cytometry. Compared with serum-free treatment, all serum-preconditioned formulations showed a clear increase in reporter expression, with transfection peaking at approximately 6% FBS and declining modestly at higher serum concentrations (Figure 6A). These results indicate that serum components can restore LNP-mediated DNA delivery, but that this effect is maximized at intermediate serum concentrations rather than exhibiting a monotonic increase in proportion to increasing serum content.

**Figure 6.**
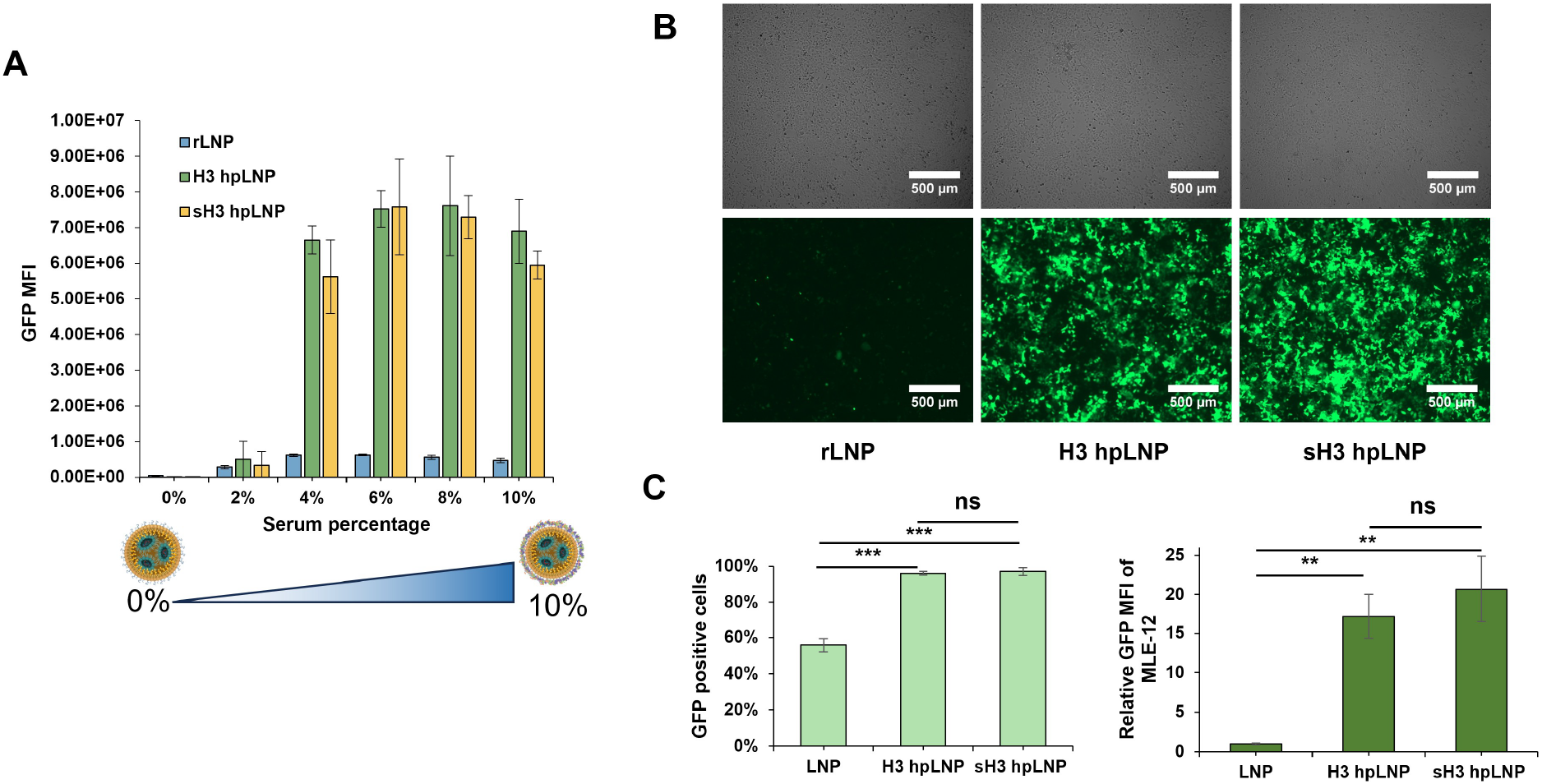
Serum-dependent gene delivery in MLE-12 cells using various GFP-encoding LNPs. (A) GFP expression analysis using flow cytometry measurements to assess transfection with various LNPs that were preconditioned at different serum percentages (v/v). Data are presented as the mean ± SD, with n=3 biological replicates. MFI = mean fluorescence intensity. (B) Representative phase contrast (top) and fluorescent (bottom) images of MLE-12 cells 48 h after transfection with various LNPs that were preconditioned at a serum percentage of 6%. (C) GFP expression levels quantified using flow cytometery analysis of MLE-12 cells treated with various LNPs following preconditioning of LNPs at a serum concentration of 6%. Data are presented as the mean ± SD, with LNPs of each type derived from different LNP batches. All MFIs were normalized to the MFI of the rLNP group. Statistical significance was calculated using a two-tailed Student’s t-test for comparisons between two sample sets. ∗∗P <0.01, ∗∗∗P < 0.001.

Peptide incorporation further enhanced transfection efficiency across all serum-conditioned formulations. For example, after conditioning in 6% FBS, hpLNPs exhibited approximately 12-fold higher GFP expression than rLNPs (Figure 6A). Notably, sH3 hpLNPs performed comparably to H3-hpLNPs, suggesting that the observed enhancement was driven by a sequence-independent effect of the peptide. Moreover, the comparable activity of H3 and sH3 hpLNPs supported a model in which peptide incorporation modifies LNP activity largely based on peptide alterations to the properties of the hpLNP surface, resulting in more favorable interactions with serum-derived components. This interpretation also is consistent with the relatively low peptide/DNA ratio used in these formulations, which may be insufficient to drive histone peptide-mediated changes to nuclear trafficking.^46^

To determine whether peptide-associated gene delivery enhancements were reproducible, we performed technical replications using three independent hpLNP batches preconditioned in 6% FBS. Across these replicates, rLNPs transfected approximately 60% of cells, whereas both H3-hpLNPs and sH3 hpLNPs consistently achieved near-complete GFP-positive populations (Figure 6B,C). Although hpLNPs showed some variation in absolute expression intensity across batches, their improvement over rLNPs remained statistically significant, reaching approximately 17-fold higher overall GFP expression without detectable loss of cell viability (Figure 6B,C and Supplemental Figure S4). These results confirmed that peptide incorporation provides a substantial and reproducible boost in LNP-mediated gene delivery while maintaining low cytotoxicity.

Finally, to evaluate whether this enhanced activity extended to non-target resident cells in the lung, we tested the gene transfection potency of the same formulations in MH-S murine alveolar macrophage cells. Because macrophages are highly phagocytic, we expected substantial nanoparticle uptake; however, we hypothesized that lysosomal processing would limit productive transgene expression. Consistent with this expectation, all LNP formulations showed minimal GFP expression in MH-S cells despite clear evidence of nanoparticle uptake (Supplemental Figure S5). These findings show that serum preconditioning enables efficient LNP-mediated DNA delivery in epithelial-like lung cells but not alveolar macrophages. Furthermore, peptide incorporation substantially amplifies this effect through a sequence-independent mechanism.

### Evaluation of serum pre-coating effect on gene transfer in murine lung

Given that serum pre-incubation markedly enhanced transfection efficiency in MLE-12 cells, we next asked whether this strategy could also enable gene transfer following local intratracheal administration in murine lung at postnatal day 7 (p7). While MLE-12 cells provide a useful AT2 cell model, they do not capture the responses of other critical lung cell types. In particular, AT1 cells play a central role in gas exchange but are terminally differentiated and therefore are difficult to evaluate through *in vitro* assessments, while alveolar macrophages may further influence delivery macroscopic outcomes in epithelial cells through nanoparticle uptake and clearance. The *in vivo* setting therefore provides an opportunity to evaluate whether serum pre-coating can support gene expression in the lung while also examining the cellular distribution of transgene signals after local administration.

Based on these considerations, we proceeded with intratracheal administration experiments in p7 mice using serum-precoated LNPs. Since our previous results showed that the transfection effect of peptide incorporation was sequence-independent, only H3-based hpLNPs were included. Prior to dosing, we confirmed the functionality of serum pre-coated LNPs following ultrafiltration and refrigerated storage. These serum-precoated particles maintained high transfection efficiency and preserved the peptide-associated boosting effect even after three weeks of storage prior to *in vivo* dosing (Supplementary Figure S6). When serum-precoated LNPs were administered intratracheally, localized GFP expression was observed in the lungs for both rLNPs and hpLNPs as compared to untreated controls, indicating that serum pre-coating augmented the activity of both LNP types and resulted in visually detectable transgene expression in vivo (Figure 7A).

**Figure 7.**
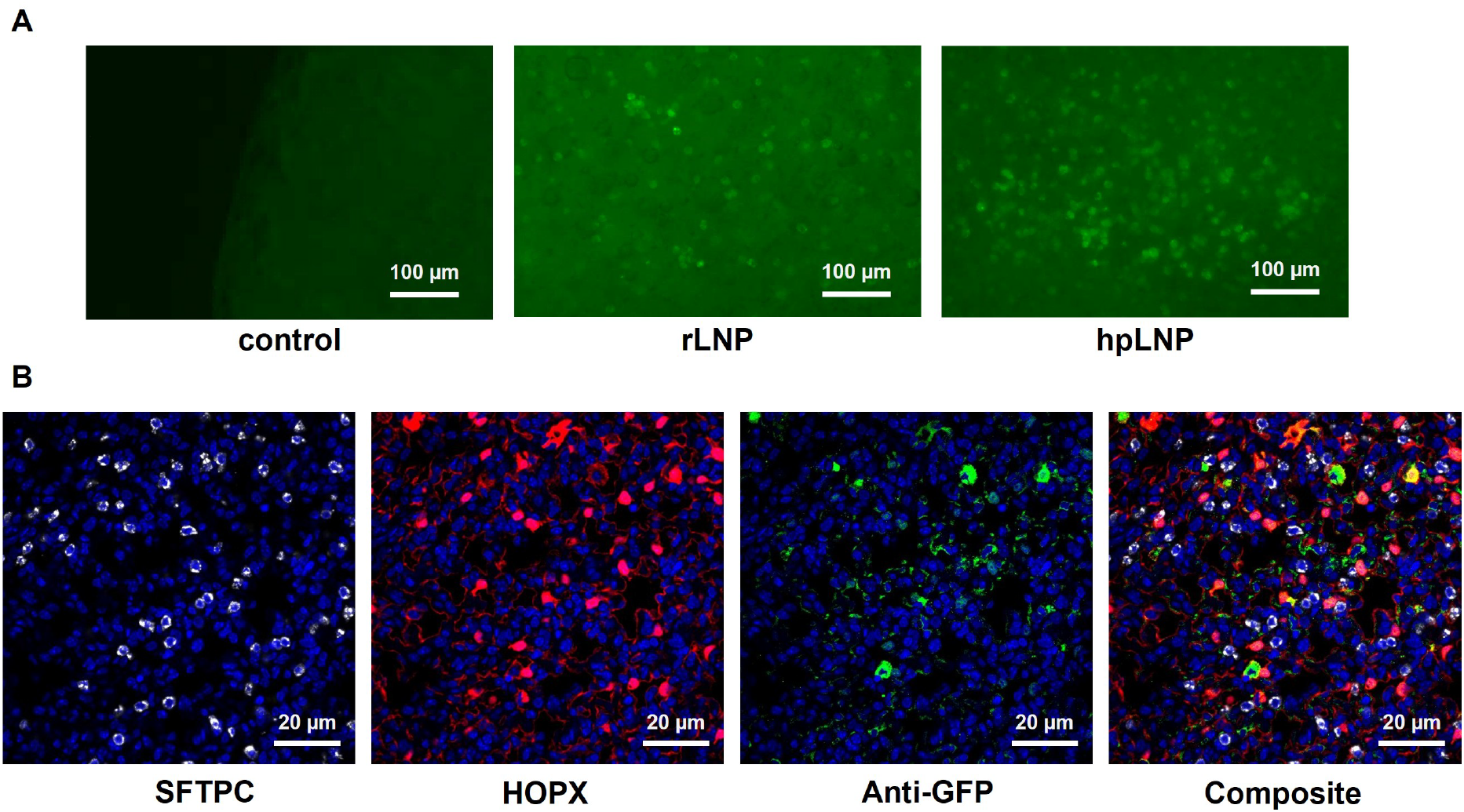
Representative fluorescent and immunohistochemistry (IHC) stained images of mouse lung 3 days post-intratrachael administration of serum pre-coated LNPs. (A) GFP expression analysis of whole lung using EVOS imaging following lung dosing with different LNPs. Control lungs were not dosed with LNPs. (B) Assessment of GFP expression colocalization with various IHC-stained markers of different epithelial cell populations in lung following dosing with serum pre-coated rLNPs.

To gain additional information regarding cellular-level gene expression resulting from administration of the serum pre-coated LNPs, we further performed immunohistochemistry on rLNP to identify which cell types in lung contained GFP. We observed that GFP expression was largely localized to AT1 cells lining the alveolar surface (stained with anti-HOPX marker), with only limited signal detected in AT2 cells (stained with anti-Surfactant Associated Protein C marker (SFTPC)), as well as some signal cells that were neither AT1 nor AT2 (Figure 7B). This distribution is consistent with the fact that AT1 cells cover ∼95% of the alveolar surface area, which may explain the relatively lower expression levels observed in AT2 cells.^47^

### Mechanism investigation of serum-conditioned hpLNPs gene expression enhancement

Since LNP transgene delivery was strongly serum-dependent and prior measurements showed minimal pKa shift, we hypothesized that enhanced expression in hpLNPs was primarily a result of peptide-mediated modulation of LNP interactions with serum components, resulting in alterations to cellular uptake. This interpretation is consistent with prior studies showing that protein corona composition can strongly influence nanoparticle uptake behavior and biological activity; for example, selective organ targeting (SORT) LNPs achieve tissue specificity through changes in surface protein adsorption induced by altered lipid charge composition.^48,49^ Accordingly, we conducted cellular uptake and nuclear accumulation experiments using Cy5-labeled plasmid DNA. Each LNP formulation, including rLNPs, H3 hpLNPs, and sH3 hpLNPs, was pre-incubated with culture media containing 0-10% FBS for 24 hours, matching the serum-exposure conditions used in the transfection assays. MLE-12 cells were treated with the Cy5-DNA– loaded LNPs for 48 h and analyzed by flow cytometry to quantify intracellular Cy5 fluorescence. Distinct serum-dependent uptake behaviors were observed across formulations. For rLNPs, Cy5 fluorescence intensity was similar across conditions with slight decrease at higher serum percentages (Figure 8A), consistent with partial inhibition of nanoparticle–cell interactions by serum protein adsorption at higher serum concentrations.^50^ In contrast, hpLNPs exhibited very low uptake in the absence of serum but showed a sharp rise in Cy5 intensity at 2% serum preconditioning, reaching maximal uptake at 6% serum preconditioning. This uptake pattern closely mirrored the transfection profile described above, supporting the idea that peptide incorporation changes how LNPs interact with serum components and, subsequently, with the cell surface. Together, these data suggest that peptide incorporation enhances transfection primarily by increasing cellular uptake.

**Figure 8.**
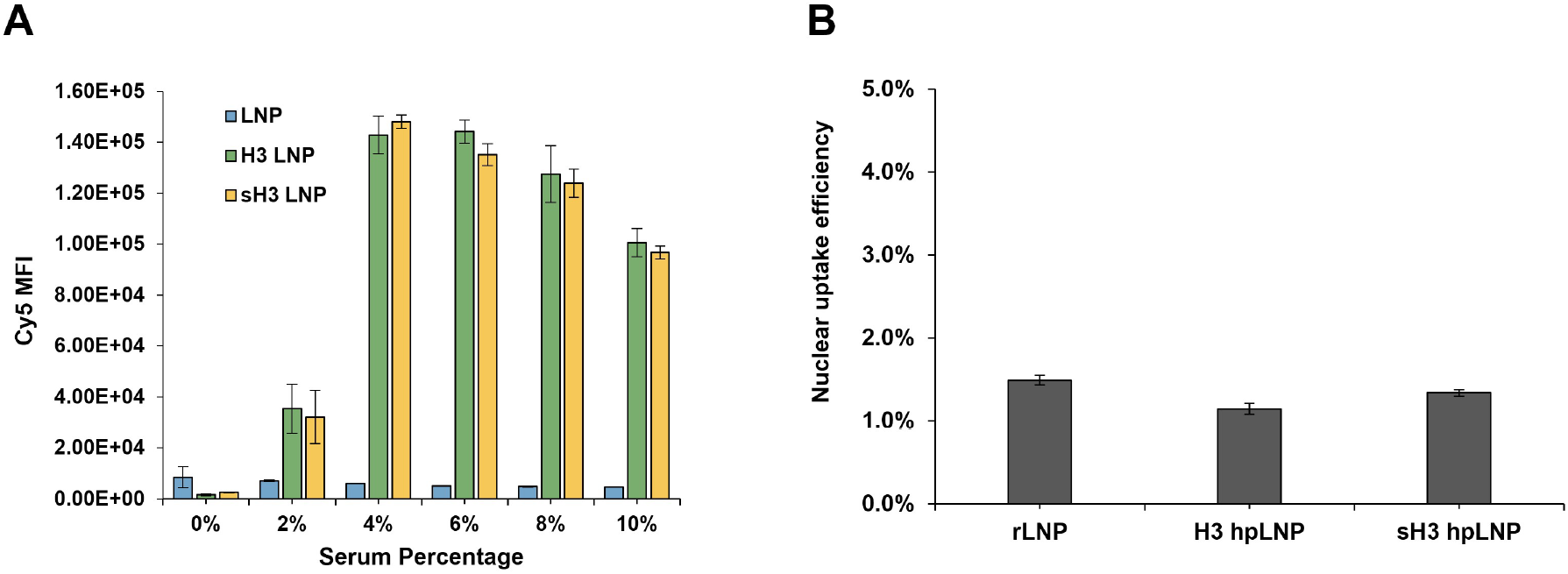
Cellular and nuclear internalization efficiency of various LNPs in MLE-12 cells based on assessment of Cy5 fluorescence. LNPs were preconditioned prior to delivery at different serum percentages. (A) Flow cytometry measurements of Cy5 intensity in MLE-12 cells 48 h after delivery of LNPs preconditioned at serum levels ranging from 0% to 10% . (B) Nuclear uptake efficiency at serum level of 6%, determined by calculating the percentage of Cy5 nuclear fluorescence relative to total Cy5 cellular fluorescence based on flow cytometry measurements. n=3 biological replicates.

We next examined nuclear uptake efficiency, to further de-couple and confirm the impacts of peptide incorporation using LNPs preconditioned at 6% serum condition. Nuclei in LNP-treated cells were isolated 48 h after transfection and analyzed via flow cytometry for nuclear Cy5 fluorescence. Although hpLNP-treated cells showed a higher absolute nuclear Cy5 intensity than rLNP-treated cells, normalization to total cell uptake revealed that the fraction of nuclear-localized DNA was comparable across all formulations (Supplemental Figure S7 and Figure 8B). This result indicated that inclusion of the peptide contributed minimally to nuclear transport. Given the relatively low peptide-to-DNA mass ratio, it is plausible that electrostatic interactions between the DNA and peptide weaken following cellular uptake, leading to dissociation prior to nuclear entry and thus diminishing the importin-mediated nuclear import capacity of the H3 peptide observed in our earlier work with H3 polyplexes.^46^

The distinct serum-dependent uptake profile of hpLNPs as compared with rLNPs suggested that peptide incorporation does not simply increase serum adsorption globally, but instead alters interactions with specific serum components. Among these components, exosomes are of particular interest because they are abundant in serum and have been reported to influence nanoparticle trafficking either through exosome-nanoparticle surface interaction or direct exosome-nanoparticle fusion.^51^ To directly test whether exosomes in serum were influencing overall uptake and gene delivery in hpLNPs, we performed transfection experiments using commercially available exosome-depleted serum. Significant depletion of exosome-sized particles in the commercially purchased FBS was confirmed by nanoparticle tracking analysis (NTA), which showed exosome concentrations in the media at levels that were consistent with the manufacturer’s specifications (Supplemental Figure S8).

When incubated with exosome-depleted FBS media, rLNPs showed a moderate increase in reporter protein expression relative to the normal serum condition, suggesting that exosome interaction may partially sequester or inhibit conventional LNPs. In contrast, hpLNPs displayed a dramatic 13-fold reduction in expression under the same exosome-depleted conditions. (Figure 9). This loss of expression when hpLNPs were preconditioned in exosome-depleted serum indicated that peptide incorporation substantially changed the interaction between LNPs and exosome-associated serum components. The reduced expression in hpLNPs without exosomes may arise from altered physicochemical interactions introduced by peptide incorporation, which could modify the internal charge distribution of the encapsulated DNA and subtly reorganize the surrounding lipid membrane. Such changes may increase the affinity of hpLNPs for serum exosome-associated components, thereby promoting hpLNP association with cells and improving uptake efficiency. Importantly, even after normalizing for the reduced total protein content in exosome-depleted serum resulting from loss of membrane-bound proteins within exosomes, hpLNPs maintained significantly lower transfection efficiency than rLNPs, confirming that exosomes play a key functional role in facilitating hpLNP uptake and gene expression (Supplemental Figure S10).

**Figure 9.**
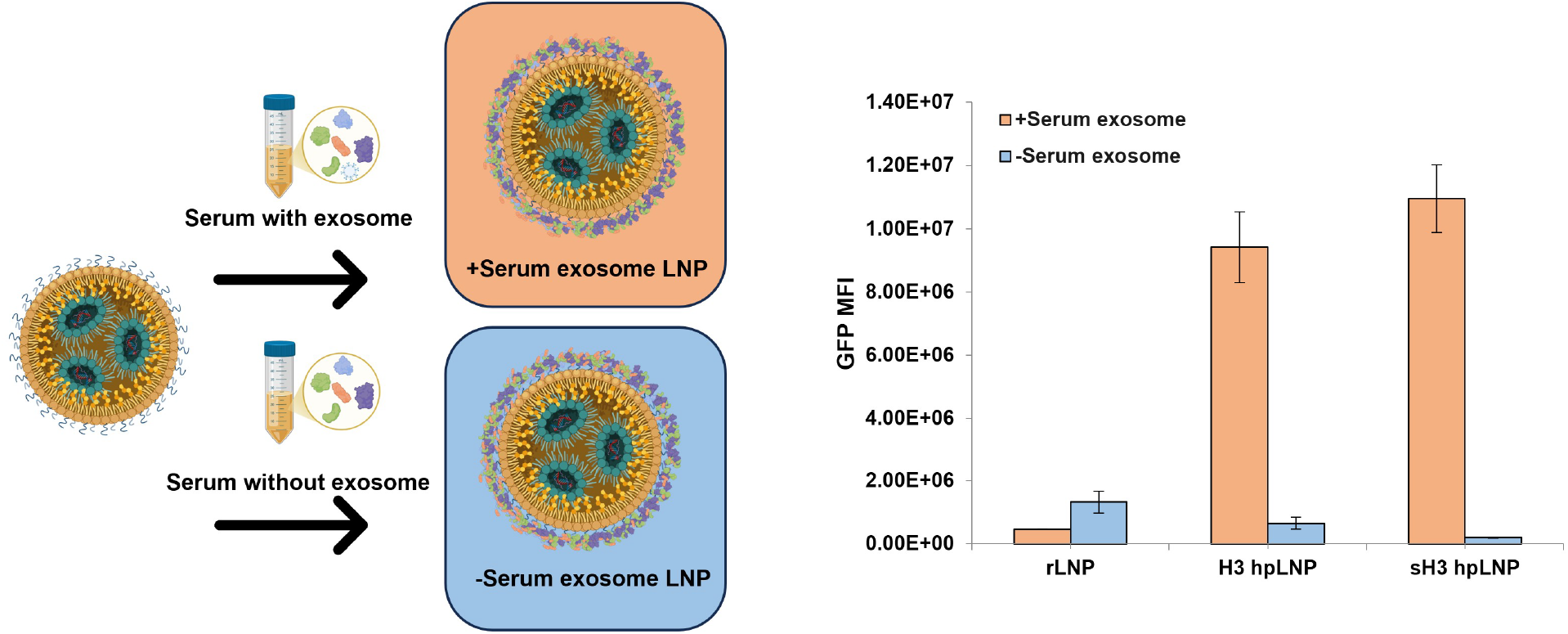
Investigation of exosome-dependent LNP transfection in MLE-12 cells. Flow cytometry measurement of GFP expression levels following transfection with LNPs that were pre-incubated with 6% serum-containing media with and without serum exosomes.

To further support the conclusion that peptide incorporation changes serum-component adsorption, we analyzed LNPs density distribution and associated protein species after serum incubation using density-gradient ultracentrifugation, a technique shown to have better separation for low density LNPs for protein corona analysis in recent year.^42^ Upon separation of fractions, we observed that all LNPs exhibited a shift toward higher-density fractions, with hpLNPs shifting down to even higher densities than the densities observed for the rLNPs, potentially due to the inherent higher density from the encapsulated peptide as well as the attached serum exosome components (Supplemental Figure S11A). When analyzing each collected peak fraction collection through protein gel electrophoresis, distinct protein peak intensities and profiles, notably at 25 and 37 kDa, were detected between rLNPs and hpLNPs (Supplemental Figure S11B and S11C). Together, the exosome-depletion studies provided strong evidence that peptide incorporation modifies LNP interactions with serum components, particularly with exosome-associated components, thereby reshaping cellular uptake behavior and accounting for the exosome-dependent enhancement of hpLNP transfection efficiency.

## Conclusion

In conclusion, short histone-inspired peptides were incorporated into a clinically-validated LNP formulation through a core-shell assembly strategy to evaluate the influence of peptide-modification in LNPs on serum component deposition and gene delivery in murine lung epithelial cells and murine lung epithelium. Under serum-free conditions, minimal expression was observed in MLE-12 cells or lung epithelium, whereas pre-conditioning LNPs or hpLNPs with serum resulted in efficient transgene expression both *in vitro* and *in vivo*. Notably, peptide inclusion in the pre-conditioned hpLNPs resulted in 17-fold higher GFP expression than in pre-conditioned rLNPs in MLE-12 cells, with the strongest overall expression observed with 6% serum pre-conditioning. Furthermore, mechanistic analyses revealed that peptide incorporation primarily altered transgene delivery by enhancing cellular uptake through alteration of serum component interactions. Under exosome-depleted conditions, hpLNP transgene expression decreased by roughly 13-fold whereas rLNP transgene expression increased, indicating that peptide additives can trigger LNP association with serum exosome components. Density gradient ultracentrifugation confirmed higher post-serum density and distinct protein profiles for hpLNPs, consistent with altered surface protein composition. Together, these findings demonstrate that peptide incorporation modulates serum component binding and enhances cellular uptake, providing a strategy for DNA delivery in serum-free environment.

## Methods

### Reagents and materials

6-((2-hexyldecanoyl)oxy)-N-(6-((2-hexyldecanoyl)oxy)hexyl)-N-(4-hydroxybutyl)hexan-1-aminium (ALC-0315), Methoxypolyethyleneglycoloxy(2000)-N,N-ditetradecylacetamide (ALC-159) and 1,2-distearoyl-sn-glycero-3-phosphocholine (DSPC) were purchased from Avanti Research (Alabaster, AL, USA). 6-(p-Toluidino)-2-naphthalenesulfonic acid sodium salt (TNS), cholesterol, and heparin were purchased from Millipore-Sigma (St. Louis, MO, USA). All peptides (H3 peptide ARTKQTARKSTGGKAPRKQLATKAA and scrambled H3 peptide LSAATPRTAKGARQTKRQKAKGTAK) were custom synthesized by Genscript at ≥95% purity (Piscataway, NJ, USA). The GWIZ-GFP mammalian expression plasmid was previously purchased from Genlantis (San Diego, CA, USA), amplified in MAX Efficiency DH5α Competent Cells from Thermo Fisher (Nazareth, PA, USA) and purified with a ZymoPURE II Plasmid Maxiprep Kit (Zymo Research, Tustin, CA, USA) in accordance with the manufacturer’s protocols. Cy5 Label IT nucleic acid labeling kit from Mirus Bio (Madison, WI, USA) was used for pDNA labeling for subsequent cellular and nucleus uptake experiments. 4′,6-diamidino-2-phenylindole (DAPI) solution was purchased from Abcam (Waltham, MA, USA). Amicon Ultra-2 Centrifugal Filter Units were purchased from Millipore-Sigma (St. Louis, MO, USA)

### hpLNP preparation

The hpLNPs were formulated via a step wise method. For the DNA peptide core, both DNA and peptide solution were dissolved in 10 mM citrate buffer (pH 4). DNA solution was keeping constant at 107 ug/ml, while adjusting peptide solutions concentration to achieve different N/P ratio (0-3.6). The N/P ratios were calculated as the ratio of the number of charged amine species (with contributions from arginine and lysine residues) to the number of phosphates in the pDNA. Subsequently, equal volumes of DNA and peptide solution were combined via pipette mixing followed by gentle vortexing. The complex was allowed to stabilize at room temperature for 30 minutes. The final LNPs were prepared by rapid mixing lipid solutions dissolved in 100% ethanol and citrate solutions containing either pDNA alone or the peptide-DNA complex using a NanoAssemblr Ignite microfluidic platform (Precision NanoSystems, Vancouver, BC, Canada). The lipid compositions used consisted of ionizable lipid ALC-0315/PEG lipid ALC-0159/Cholesterol/DSPC at a molar ratio of 47.5:1.8:40.7:10 and a total lipid molar concentration of 20 mM, equivalent to a final weight ratio of 40.2:1 (total lipids/DNA).^37^ The total flow rate for the mixing was 12 mL/min, with flow rate ratio of 3:1 for the aqueous to ethanol stream. Immediately after mixing, the crude LNP solution was transferred to a 3.5 kDa MWCO Pur-A-Lyzer Maxi Kit (Sigma-Aldrich, Milwaukee, WI, USA) and serially dialyzed two times against 1X phosphate-buffered saline (PBS) pH 7.4 at 1:250 (v:v) for two h each time. The final neutral LNP solutions were stored at 4 °C until use.

### rLNP and hpLNP characterization

The size and the polydispersity index (PDI) for both the rLNPs and hpLNPs were measured with a Zetasizer Advance instrument (Malvern Panalytical, Malvern, UK). Samples were diluted 10X in PBS and loaded onto a low volume cuvette for measurements. For nanoparticle serum stability testing, LNPs containing 1 μg pDNA were added into wells containing serum percentages ranging from 0-10% FBS and the resulting mixed samples were placed in a 37°C incubator. Size measurements were taken at 0, 7, 16, 30, 44 h post incubation. For nanoparticle storage stability assessments, size measurements were taken on the day of nanoparticle synthesis and 6 months post synthesis following storage at 4°C.

Encapsulation was confirmed via both DNA gel electrophoresis and Quant-iT PicoGreen assays (Thermo Fisher) following the manufacturer’s protocol. Equal amounts of DNA-loaded LNPs were analyzed on a 1% agarose gel containing SYBR Safe DNA dye under three conditions: treatment with 0.2% Triton X-100 in PBS to release encapsulated cargo, treatment with 0.6 mg/ml heparin to release DNA from peptide interactions, or treatment with both additives in combination. After treatment, the mixtures were incubated at 37°C for 10 minutes. The mixtures were then loaded onto the agarose gel with glycerol loading buffer, run at 100 V for 1 hour, and imaged with a Bio-Rad GelDoc Go imager (Hercules, CA, USA). The same treated mixtures were used to determine the encapsulation efficiency of pDNA. The relative values of free mRNA and encapsulated mRNA were calculated based on fluorescence intensity from PicoGreen using the following formula:

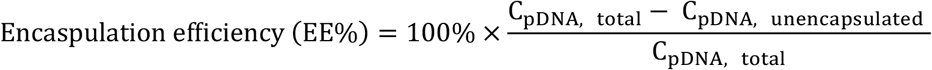

Where each of the pDNA concentrations were calculated from PicoGreen assay corresponding standard curve. The resulting encapsulated DNA concentrations were used as effective concentrations for further in vitro and in vivo dosing.

The apparent pKa value of each type of LNP was determined via a TNS assay adapted from a previously described protocol.^52^ Buffer solutions ranging from 3 to 10 were prepared by titrating 20 mM sodium citrate dihydrate/citric acid (containing 150 mM sodium chloride, pH 3-5.5, in 0.5 pH increments), 20 mM dipotassium phosphate/monopotassium phosphate (containing 150 mM sodium chloride, pH 6-8, in 0.5 pH increments), and 20 mM Tris/HCl (containing 150 mM sodium chloride, pH 8.5-10, in 0.5 pH increments). A 0.6 mM stock TNS solution was prepared in biograde water. In a 96-well black plate, 2 μl of TNS solution and 20 μl of each LNP solution (Diluted to same concentration) were added into 178 μl of corresponding gradient pH buffer (with a total of 16 different pH points) and mixed thoroughly. After 10 minutes, the fluorescence intensity of the TNS was measured using BioTek Synergy H4 plate reader (Winooski, VT, USA), and the half point selected from a sigmoid fit was calculated to be the apparent pKa of each LNP.

Cryo-TEM images were acquired using a Thermo Scientific Talos L120C transmission electron microscope. Prior to imaging, 3.5 μl of sample was pipetted onto a freshly glow-discharged 300 mesh copper lacey carbon grid and plunge frozen into liquid ethane using a Thermo Scientific Vitrobot Mark IV maintained at a temperature of 5°C and a relative humidity of 100%. Vitrified samples were maintained at -170°C and imaged in low-dose mode using the Thermo Scientific EPU software. Images were collected at 120 kV with a Ceta 16M camera using a total dose of 30 electrons/Å^2^ and defocus values of -3.0, -3.5, and -4 μm.

### *In vitro* MLE-12 cell transfection assays

Murine MLE-12 cells were purchased from American Type Culture Collection (ATCC, Manassas, VA, USA) and cultured in Corning Dulbecco’s Modified Eagle’s Medium (DMEM)/Hams F-12 50/50 Mix containing 10% heat inactivated fetal bovine serum (FBS) and 100 U/mL of penicillin/streptomycin, maintained at 37°C and 5% carbon dioxide. MH-S cells were cultured in RPMI-1640 containing 10% FBS, 1% pen/strep and 0.05 mM 2-mercaptoethanol, and also maintained at 37°C and 5% carbon dioxide.

To evaluate transfection, MLE-12 cells were seeded and grown in a 24-well plate at a seeding 20,000 cells/cm^2^ for 24 h before the experiments. In parallel, LNP samples containing 1 μg pDNA were added into growth media in the presence of varying quantities of serum (0-10% serum/media by volume) and briefly mixed before pre-incubation at 37°C for 24 h to ensure a stable protein corona formation. After 24 h, pre-incubated LNPs were added to cells by replacing the culture media with the media containing pre-incubated LNPs. Cells were imaged with a Zeiss Axio 7 Observer (White Plains, NY) at 24 and 48 h post transfection. After two days of incubation, the cells were washed with DPBS and prepared for flow cytometry measurements.

GFP expression and Cy5 intensity as a measure of cell uptake were quantified using a Novocyte Flow Cytometer (Agilent Technologies). For flow cytometry, cells were detached after image using a standard trypsin-based protocol: 200 μl of 0.25% of trypsin was added into each well and incubated for 5 minutes for cell detachment, and subsequently, a flow solution (Dulbecco’s PBS with 2% FBS) was added to neutralize the enzymatic process. The cells were washed two more times with flow solution and transferred into flow tubes for flow analysis. A total of 30,000 cells events were collected in each sample for analysis of each well. For cell uptake analysis, cells were washed with CellScrub Buffer (Amsbio, Cambridge, MA, USA) for 10 mins prior to standard flow sample preparation to remove any extracellularly attached LNPs.

For nuclear uptake analysis, one half of the Cy5 LNP-dosed cells were prepared for nuclei isolation. Immediately after trypsin detachment and flow sample preparation, cell membranes were lysed with Nuclei Isolation Lysis Reagent (NILR) using a procedure adapted from the Takara Bio nuclei isolation protocol.^53^ The NILR consists of 1:100 4, 6-diamidino-2-phenylindole (DAPI) solution, 1% (v/v) of 1M Tris-HCl and NaCl solution, 0.3% (v/v) of 1M MgCl_2_ solution, and 1% (v/v) non-ionic surfactant TERGITOL solution (Millipore-Sigma, Burlington, MA) in water. After 10 minutes of incubation with the NILR, the lysed cell solution was centrifuged at 300 g for 5 minutes and resuspended in flow solution. The nuclear uptake efficiency was calculated using the formula below, where the mean fluorescent intensity (MFI) of both the nuclear and whole cell fractions were used.

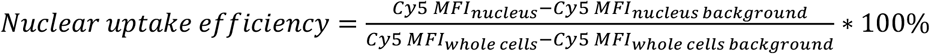

For exosome-dependent cell transfections, exosome-depleted FBS obtained from Thermo Fisher (Nazareth, PA, USA) was used for media supplementation instead of regular FBS. The number and the size of particles in both of types of serum were measured using a nanoparticle tracking analysis (NTA) device ZetaView (Particle Metrix, Germany).

### Gradient protein corona isolation

The LNPs with Cy5 DNA or a PBS control were incubated for 24h at 37°C with media containing 6% FBS and the solution was subsequently concentrated back to its original volume using a 100 kDa Amicon ultra centrifugal filter (Millipore Sigma, Burlington, MA). A density gradient protein corona isolation adapted from a previously described protocol was used.^42^ A total of 400 μl of pre-incubated solution was combined with an equal volume of 60% iodixanol Optiprep density gradient medium (Progen Biotech Inc, Wayne, PA) to achieve a final 30% iodixanol solution by volume. This mixture was subsequently loaded onto the bottom of a 5 mL ultracentrifuge tube. Next, 5 additional iodixanol layers (840 μl per layer) with 5% decrements in density (25%, 20%, 15%, 10%, and 5%) were carefully loaded above the bottom layer containing LNPs. Samples were then centrifuged for 16.5 h at 160,000g and 4°C with minimum acceleration and no braking in a SW55 Ti Beckman swinging bucket rotor. 200 μl of the mixture at a time was carefully removed and added into a black 96-well plate. A total of 24 fractions were analyzed per sample. The Cy5 fluorescence of each fraction was measured in a BioTek Synergy H4 plate reader (Winooski, VT, USA) to determine the location of the LNP in each layer. The peak fractions containing LNP were pooled and further concentrated using a 3 kDa Amicon® Ultra Centrifugal Filter for SDS-PAGE protein gel analysis.

For protein gel analysis, each fraction was diluted to an equal 30 μl volume and processed through standard SDS-PAGE sample preparation. First, 10 μl of 4X SDS loading buffer was added into the selected sample fraction prior to denaturing at 95°C for 30 mins. Next, 10 μl of each sample was loaded onto a gradient 4– 20% precast polyacrylamide gel from Bio-Rad (Hercules, CA, USA) and separated by electrophoresis at 250 V for 30 mins. The final gel was stained with SYPRO protein gel stain from Thermo Fisher (Nazareth, PA, USA) and imaged in Bio-Rad GelDoc Go Gel Imaging System.

### Intra-tracheal delivery and analysis

Prior to *in vivo* dosing, rLNP and H3 hpLNPs were pre-incubated in media containing 0% or 6 % FBS for 24 hours. Subsequently, each LNPs were concentrated down to 80 ug/ml of effective DNA concentration using 100 kDa Amicon ultra centrifugal filters. Seven-day old WT or GFP positive CD1 mouse pups were used for intra-tracheal administration of LNPs. Initially, 3% isoflurane was used to induce and maintain anesthesia of the pups throughout the minor non-invasive surgical procedure. The pups were placed on dorsal recumbency on a heated pad, and an engineered nose cone designed to fit the pup nostrils was used to maintain the flow of oxygen and anesthetic agent. The ventral neck region was prepared aseptically, and 1 cm skin incision was made midway along the neck, followed by a blunt dissection using a pair of forceps to reveal the trachea. A 50 μL Neuros Hamilton syringe fitted with a 33-gauge needle was used to administer LNPs directly into the trachea just posterior to the larynx. Pups were randomized to receive 15 μl of desired nanoparticle or control. All the animal studies were conducted at Nemours Biomedical Research Center and were approved by IACUC at Nemours. Animals were euthanized 3 days after intra-tracheal instillation. Both lungs were removed en-bloc and spatial distribution for GFP was assessed by live imaging). The lungs were then fixed in 2% paraformaldehyde followed by serial dehydration and embedded in paraffin for immunohistochemical (IHC) analysis. IHC to detect proteins was performed using the following antibodies on paraffin sections: GFP (chicken, 1:500; Aves), SFTPC (rabbit, 1:250; Abcam), and HOPX (mouse, 1:50; Santa Cruz Biotechnology).

## Author contribution statement

**Jinzhen Hu:** Conceptualization, Methodology, Investigation, Data curation, Validation, Formal analysis, Writing-original draft, Writing - review & editing; **Michael B. Papah:** Methodology, Investigation, Data curation, Validation, Formal analysis, Writing - review & editing; **Arlett Ramirez:** Methodology, Investigation, Data curation, Validation, Formal analysis, Writing - review & editing; **Deepthi Alapati:** Conceptualization, Writing - review & editing, Supervision, Resources, Funding acquisition; **Millicent O. Sullivan:** Conceptualization, Writing - review & editing, Supervision, Resources, Funding acquisition.

## Acknowledgements

The authors would like to thank Shannon Modla from Bioimaging Center in Delaware Biotechnology Institute for collecting the cryogenic TEM of all types of LNPs. The draft of the manuscript was edited using ChatGPT and OpenAI to polish grammar and assist in organizing manuscript sections in the initial draft. BioRender software was used for creating figures and schematics.

## Note

The authors declare no competing financial interest.

## References

(1) Barbier, A. J.; Jiang, A. Y.; Zhang, P.; Wooster, R.; Anderson, D. G. The Clinical Progress of mRNA Vaccines and Immunotherapies. Nat. Biotechnol. 2022, 40 (6), 840–854. 10.1038/s41587-022-01294-2.

(2) Parhiz, H.; Atochina-Vasserman, E. N.; Weissman, D. mRNA-Based Therapeutics: Looking beyond COVID-19 Vaccines. The Lancet 2024, 403 (10432), 1192–1204. 10.1016/S0140-6736(23)02444-3.

(3) Cao, J.; An, D.; Galduroz, M.; Zhuo, J.; Liang, S.; Eybye, M.; Frassetto, A.; Kuroda, E.; Funahashi, A.; Santana, J.; Mihai, C.; Benenato, K. E.; Kumarasinghe, E. S.; Sabnis, S.; Salerno, T.; Coughlan, K.; Miracco, E. J.; Levy, B.; Besin, G.; Schultz, J.; Lukacs, C.; Guey, L.; Finn, P.; Furukawa, T.; Giangrande, P. H.; Saheki, T.; Martini, P. G. V. mRNA Therapy Improves Metabolic and Behavioral Abnormalities in a Murine Model of Citrin Deficiency. Mol. Ther. 2019, 27 (7), 1242–1251. 10.1016/j.ymthe.2019.04.017.

(4) Yamazaki, K.; Kubara, K.; Ishii, S.; Kondo, K.; Suzuki, Y.; Miyazaki, T.; Mitsuhashi, K.; Ito, M.; Tsukahara, K. Lipid Nanoparticle-Targeted mRNA Formulation as a Treatment for Ornithine-Transcarbamylase Deficiency Model Mice. Mol. Ther. -Nucleic Acids 2023, 33, 210–226. 10.1016/j.omtn.2023.06.023.

(5) Zhu, X.; Yin, L.; Theisen, M.; Zhuo, J.; Siddiqui, S.; Levy, B.; Presnyak, V.; Frassetto, A.; Milton, J.; Salerno, T.; Benenato, K. E.; Milano, J.; Lynn, A.; Sabnis, S.; Burke, K.; Besin, G.; Lukacs, C. M.; Guey, L. T.; Finn, P. F.; Martini, P. G. V. Systemic mRNA Therapy for the Treatment of Fabry Disease: Preclinical Studies in Wild-Type Mice, Fabry Mouse Model, and Wild-Type Non-Human Primates. Am. J. Hum. Genet. 2019, 104 (4), 625–637. 10.1016/j.ajhg.2019.02.003.

(6) Rojas, L. A.; Sethna, Z.; Soares, K. C.; Olcese, C.; Pang, N.; Patterson, E.; Lihm, J.; Ceglia, N.; Guasp, P.; Chu, A.; Yu, R.; Chandra, A. K.; Waters, T.; Ruan, J.; Amisaki, M.; Zebboudj, A.; Odgerel, Z.; Payne, G.; Derhovanessian, E.; Müller, F.; Rhee, I.; Yadav, M.; Dobrin, A.; Sadelain, M.; Łuksza, M.; Cohen, N.; Tang, L.; Basturk, O.; Gönen, M.; Katz, S.; Do, R. K.; Epstein, A. S.; Momtaz, P.; Park, W.; Sugarman, R.; Varghese, A. M.; Won, E.; Desai, A.; Wei, A. C.; D’Angelica, M. I.; Kingham, T. P.; Mellman, I.; Merghoub, T.; Wolchok, J. D.; Sahin, U.; Türeci, Ö.; Greenbaum, B. D.; Jarnagin, W. R.; Drebin, J.; O’Reilly, E. M.; Balachandran, V. P. Personalized RNA Neoantigen Vaccines Stimulate T Cells in Pancreatic Cancer. Nature 2023, 618 (7963), 144–150. 10.1038/s41586-023-06063-y.

(7) Hamouda, A. E. I.; Filtjens, J.; Brabants, E.; Kancheva, D.; Debraekeleer, A.; Brughmans, J.; Jacobs, L.; Bardet, P. M. R.; Knetemann, E.; Lefesvre, P.; Allonsius, L.; Gontsarik, M.; Varela, I.; Crabbé, M.; Clappaert, E. J.; Cappellesso, F.; Caro, A. A.; Gordún Peiró, A.; Fredericq, L.; Hadadi, E.; Estapé Senti, M.; Schiffelers, R.; van Grunsven, L. A.; Aboubakar Nana, F.; De Geest, B. G.; Deschoemaeker, S.; De Koker, S.; Lambolez, F.; Laoui, D. Intratumoral Delivery of Lipid Nanoparticle-Formulated mRNA Encoding IL-21, IL-7, and 4-1BBL Induces Systemic Anti-Tumor Immunity. Nat. Commun. 2024, 15 (1), 10635. 10.1038/s41467-024-54877-9.

(8) Fu, Q.; Zhao, X.; Hu, J.; Jiao, Y.; Yan, Y.; Pan, X.; Wang, X.; Jiao, F. mRNA Vaccines in the Context of Cancer Treatment: From Concept to Application. J. Transl. Med. 2025, 23 (1), 12. 10.1186/s12967-024-06033-6.

(9) Olson, K. E.; Namminga, K. L.; Lu, Y.; Thurston, M. J.; Schwab, A. D.; de Picciotto, S.; Tse, S.-W.; Walker, W.; Iacovelli, J.; Small, C.; Wipke, B. T.; Mosley, R. L.; Huang, E.; Gendelman, H. E. Granulocyte-Macrophage Colony-Stimulating Factor mRNA and Neuroprotective Immunity in Parkinson’s Disease. Biomaterials 2021, 272, 120786. 10.1016/j.biomaterials.2021.120786.

(10) Dong, H.; He, Z.; Cai, S.; Ma, H.; Su, L.; Li, J.; Yang, H.; Xie, R. Methylprednisolone Substituted Lipid Nanoparticles Deliver C3 Transferase mRNA for Combined Treatment of Spinal Cord Injury. J. Nanobiotechnology 2025, 23 (1), 98. 10.1186/s12951-025-03153-z.

(11) Padda, I. S.; Mahtani, A. U.; Patel, P.; Parmar, M. Small Interfering RNA (siRNA) Therapy. In StatPearls; StatPearls Publishing: Treasure Island (FL), 2026.

(12) Zhang, W.; Pfeifle, A.; Lansdell, C.; Frahm, G.; Cecillon, J.; Tamming, L.; Gravel, C.; Gao, J.; Thulasi Raman, S. N.; Wang, L.; Sauve, S.; Rosu-Myles, M.; Li, X.; Johnston, M. J. W. The Expression Kinetics and Immunogenicity of Lipid Nanoparticles Delivering Plasmid DNA and mRNA in Mice. Vaccines 2023, 11 (10), 1580. 10.3390/vaccines11101580.

(13) Lu, B.; Lim, J. M.; Yu, B.; Song, S.; Neeli, P.; Sobhani, N.; K P.; Bonam, S. R.; Kurapati, R.; Zheng, J.; Chai, D. The Next-Generation DNA Vaccine Platforms and Delivery Systems: Advances, Challenges and Prospects. Front. Immunol. 2024, 15. 10.3389/fimmu.2024.1332939.

(14) Neeli, P.; Chai, D.; Roy, D.; Prajapati, S.; Bonam, S. R. DNA Vaccines in the Post-mRNA Era: Engineering, Applications, and Emerging Innovations. Int. J. Mol. Sci. 2025, 26 (17), 8716. 10.3390/ijms26178716.

(15) Packer, M.; Gyawali, D.; Yerabolu, R.; Schariter, J.; White, P. A Novel Mechanism for the Loss of mRNA Activity in Lipid Nanoparticle Delivery Systems. Nat. Commun. 2021, 12 (1), 6777. 10.1038/s41467-021-26926-0.

(16) Cheng, F.; Wang, Y.; Bai, Y.; Liang, Z.; Mao, Q.; Liu, D.; Wu, X.; Xu, M. Research Advances on the Stability of mRNA Vaccines. Viruses 2023, 15 (3), 668. 10.3390/v15030668.

(17) Kozak, M.; Hu, J. DNA Vaccines: Their Formulations, Engineering and Delivery. Vaccines 2024, 12 (1), 71. 10.3390/vaccines12010071.

(18) Maslow, J. N.; Kwon, I.; Kudchodkar, S. B.; Kane, D.; Tadesse, A.; Lee, H.; Park, Y. K.; Muthumani, K.; Roberts, C. C. DNA Vaccines for Epidemic Preparedness: SARS-CoV-2 and Beyond. Vaccines 2023, 11 (6), 1016. 10.3390/vaccines11061016.

(19) Bozza, M.; Green, E. W.; Espinet, E.; Roia, A. D.; Klein, C.; Vogel, V.; Offringa, R.; Williams, J. A.; Sprick, M.; Harbottle, R. P. Novel Non-Integrating DNA Nano-S/MAR Vectors Restore Gene Function in Isogenic Patient-Derived Pancreatic Tumor Models. Mol. Ther. Methods Clin. Dev. 2020, 17, 957–968. 10.1016/j.omtm.2020.04.017.

(20) Bozza, M.; De Roia, A.; Correia, M. P.; Berger, A.; Tuch, A.; Schmidt, A.; Zörnig, I.; Jäger, D.; Schmidt, P.; Harbottle, R. P. A Nonviral, Nonintegrating DNA Nanovector Platform for the Safe, Rapid, and Persistent Manufacture of Recombinant T Cells. Sci. Adv. 2021, 7 (16), eabf1333. 10.1126/sciadv.abf1333.

(21) Patel, M. N.; Tiwari, S.; Wang, Y.; O’Neill, S.; Wu, J.; Omo-Lamai, S.; Espy, C.; Chase, L. S.; Majumder, A.; Hoffman, E.; Shah, A.; Sárközy, A.; Katzen, J.; Pardi, N.; Brenner, J. S. Safer Non-Viral DNA Delivery Using Lipid Nanoparticles Loaded with Endogenous Anti-Inflammatory Lipids. Nat. Biotechnol. 2026, 44 (1), 79–89. 10.1038/s41587-025-02556-5.

(22) Guillot, L.; Nathan, N.; Tabary, O.; Thouvenin, G.; Le Rouzic, P.; Corvol, H.; Amselem, S.; Clement, A. Alveolar Epithelial Cells: Master Regulators of Lung Homeostasis. Int. J. Biochem. Cell Biol. 2013, 45 (11), 2568–2573. 10.1016/j.biocel.2013.08.009.

(23) Mason, R. J. Biology of Alveolar Type II Cells. Respirology 2006, 11 (1), S12–S15. 10.1111/j.1440-1843.2006.00800.x.

(24) Han, S.; Budinger, G. R. S.; Gottardi, C. J. Alveolar Epithelial Regeneration in the Aging Lung. J. Clin. Invest. 2023, 133 (20). 10.1172/JCI170504.

(25) Choi, H. K.; Bang, G.; Shin, J. H.; Shin, M. H.; Woo, A.; Kim, S. Y.; Lee, S. H.; Kim, E. Y.; Shim, H. S.; Suh, Y. J.; Kim, H. E.; Lee, J. G.; Choi, J.; Lee, J. H.; Kim, C. H.; Park, M. S. Regenerative Capacity of Alveolar Type 2 Cells Is Proportionally Reduced Following Disease Progression in Idiopathic Pulmonary Fibrosis-Derived Organoid Cultures. Tuberc. Respir. Dis. 2024, 88 (1), 130–137. 10.4046/trd.2024.0094.

(26) Olajuyin, A. M.; Zhang, X.; Ji, H.-L. Alveolar Type 2 Progenitor Cells for Lung Injury Repair. Cell Death Discov. 2019, 5 (1), 63. 10.1038/s41420-019-0147-9.

(27) Zacharias, W. J.; Frank, D. B.; Zepp, J. A.; Morley, M. P.; Alkhaleel, F. A.; Kong, J.; Zhou, S.; Cantu, E.; Morrisey, E. E. Regeneration of the Lung Alveolus by an Evolutionarily Conserved Epithelial Progenitor. Nature 2018, 555 (7695), 251–255. 10.1038/nature25786.

(28) Nureki, S.-I.; Tomer, Y.; Venosa, A.; Katzen, J.; Russo, S. J.; Jamil, S.; Barrett, M.; Nguyen, V.; Kopp, M.; Mulugeta, S.; Beers, M. F. Expression of Mutant Sftpc in Murine Alveolar Epithelia Drives Spontaneous Lung Fibrosis. J. Clin. Invest. 2018, 128 (9), 4008–4024. 10.1172/JCI99287.

(29) Parimon, T.; Yao, C.; Stripp, B. R.; Noble, P. W.; Chen, P. Alveolar Epithelial Type II Cells as Drivers of Lung Fibrosis in Idiopathic Pulmonary Fibrosis. Int. J. Mol. Sci. 2020, 21 (7), 2269. 10.3390/ijms21072269.

(30) Guan, X.; Pei, Y.; Song, J. DNA-Based Nonviral Gene Therapy─Challenging but Promising. Mol. Pharm. 2024, 21 (2), 427–453. 10.1021/acs.molpharmaceut.3c00907.

(31) Munye, M. M.; Tagalakis, A. D.; Barnes, J. L.; Brown, R. E.; McAnulty, R. J.; Howe, S. J.; Hart, S. L. Minicircle DNA Provides Enhanced and Prolonged Transgene Expression Following Airway Gene Transfer. Sci. Rep. 2016, 6 (1), 23125. 10.1038/srep23125.

(32) van Straten, D.; Sork, H.; van de Schepop, L.; Frunt, R.; Ezzat, K.; Schiffelers, R. M. Biofluid Specific Protein Coronas Affect Lipid Nanoparticle Behavior in Vitro. J. Controlled Release 2024, 373, 481–492. 10.1016/j.jconrel.2024.07.044.

(33) Dilliard, S. A.; Cheng, Q.; Siegwart, D. J. On the Mechanism of Tissue-Specific mRNA Delivery by Selective Organ Targeting Nanoparticles. Proc. Natl. Acad. Sci. 2021, 118 (52), e2109256118. 10.1073/pnas.2109256118.

(34) Larsen, J. D.; Reilly, M. J.; Sullivan, M. O. Using the Epigenetic Code To Promote the Unpackaging and Transcriptional Activation of DNA Polyplexes for Gene Delivery. Mol. Pharm. 2012, 9 (5), 1041–1051. 10.1021/mp200373p.

(35) Larsen, J. D.; Ross, N. L.; Sullivan, M. O. Requirements for the Nuclear Entry of Polyplexes and Nanoparticles during Mitosis: Nuclear Entry during Mitosis. J. Gene Med. 2012, 14 (9–10), 580–589. 10.1002/jgm.2669.

(36) Reilly, M. J.; Larsen, J. D.; Sullivan, M. O. Histone H3 Tail Peptides and Poly(Ethylenimine) Have Synergistic Effects for Gene Delivery. Mol. Pharm. 2012, 9 (5), 1031–1040. 10.1021/mp200372s.

(37) Nauta, M.; Peeters, D. J.; Doorslaer, T. F. S. V.; Badkar, A. V.; Darvari, R.; Warne, N. W.; Jean, J.; Hendrikse, D. P. G.; Sahin, U.; GÜLER, A.; Kuhn, A.; Muik, A.; Vogel, A.; Walzer, K.; Witzel, S.; Hein, S.; TÜreci, Ö. Coronavirus Vaccine. WO2021213945A1, October 28, 2021. https://patents.google.com/patent/WO2021213945A1/en (accessed 2023-05-28).

(38) Kim, B.; Park, C. H.; Jung, I.-Y.; Lee, Y.; Lim, S. G.; Lee, D.; Kwon, S. J.; Koo, H. Size Control of Lipid Nanoparticles via Simulation-Based Design of a Microfluidic Chip and Its Effect on mRNA Delivery in Vitro and in Vivo. J. Nanobiotechnology 2025, 23 (1), 797. 10.1186/s12951-025-03836-7.

(39) Wang, R.; He, J.; Xu, Y.; Peng, B. Impact of Protein Coronas on Lipid Nanoparticle Uptake and Endocytic Pathways in Cells. Molecules 2024, 29 (20), 4818. 10.3390/molecules29204818.

(40) Straten, D. van; Schepop, L. van de; Frunt, R.; Vader, P.; Schiffelers, R. M. Serum Heat Inactivation Diminishes ApoE-Mediated Uptake of D-Lin-MC3-DMA Lipid Nanoparticles. Beilstein J. Nanotechnol. 2025, 16 (1), 740–748. 10.3762/bjnano.16.57.

(41) Berger, M.; Degey, M.; Leblond Chain, J.; Maquoi, E.; Evrard, B.; Lechanteur, A.; Piel, G. Effect of PEG Anchor and Serum on Lipid Nanoparticles: Development of a Nanoparticles Tracking Method. Pharmaceutics 2023, 15 (2), 597. 10.3390/pharmaceutics15020597.

(42) Voke, E.; Arral, M. L.; Squire, H. J.; Lin, T.-J.; Zheng, L.; Coreas, R.; Lui, A.; Iavarone, A. T.; Pinals, R. L.; Whitehead, K. A.; Landry, M. P. Protein Corona Formed on Lipid Nanoparticles Compromises Delivery Efficiency of mRNA Cargo. Nat. Commun. 2025, 16 (1), 8699. 10.1038/s41467-025-63726-2.

(43) Aliakbarinodehi, N.; Gallud, A.; Mapar, M.; Wesén, E.; Heydari, S.; Jing, Y.; Emilsson, G.; Liu, K.; Sabirsh, A.; Zhdanov, V. P.; Lindfors, L.; Esbjörner, E. K.; Höök, F. Interaction Kinetics of Individual mRNA-Containing Lipid Nanoparticles with an Endosomal Membrane Mimic: Dependence on pH, Protein Corona Formation, and Lipoprotein Depletion. ACS Nano 2022, 16 (12), 20163–20173. 10.1021/acsnano.2c04829.

(44) Ogris, M.; Brunner, S.; Schüller, S.; Kircheis, R.; Wagner, E. PEGylated DNA/Transferrin–PEI Complexes: Reduced Interaction with Blood Components, Extended Circulation in Blood and Potential for Systemic Gene Delivery. Gene Ther. 1999, 6 (4), 595–605. 10.1038/sj.gt.3300900.

(45) Gwak, S.-J.; Macks, C.; Bae, S.; Cecil, N.; Lee, J. S. Physicochemical Stability and Transfection Efficiency of Cationic Amphiphilic Copolymer/pDNA Polyplexes for Spinal Cord Injury Repair. Sci. Rep. 2017, 7 (1), 11247. 10.1038/s41598-017-10982-y.

(46) Ross, N. L.; Sullivan, M. O. Importin-4 Regulates Gene Delivery by Enhancing Nuclear Retention and Chromatin Deposition by Polyplexes. Mol. Pharm. 2015, 12 (12), 4488–4497. 10.1021/acs.molpharmaceut.5b00645.

(47) Yang, J.; Hernandez, B. J.; Martinez Alanis, D.; Narvaez del Pilar, O.; Vila-Ellis, L.; Akiyama, H.; Evans, S. E.; Ostrin, E. J.; Chen, J. The Development and Plasticity of Alveolar Type 1 Cells. Development 2016, 143 (1), 54–65. 10.1242/dev.130005.

(48) Cheng, Q.; Wei, T.; Farbiak, L.; Johnson, L. T.; Dilliard, S. A.; Siegwart, D. J. Selective Organ Targeting (SORT) Nanoparticles for Tissue-Specific mRNA Delivery and CRISPR–Cas Gene Editing. Nat. Nanotechnol. 2020, 15 (4), 313–320. 10.1038/s41565-020-0669-6.

(49) Dilliard, S. A.; Cheng, Q.; Siegwart, D. J. On the Mechanism of Tissue-Specific mRNA Delivery by Selective Organ Targeting Nanoparticles. Proc. Natl. Acad. Sci. U. S. A. 2021, 118 (52), e2109256118. 10.1073/pnas.2109256118.

(50) Bates, S. M.; Munson, M. J.; Trovisco, V.; Pereira, S.; Miller, S. R.; Sabirsh, A.; Betts, C. J.; Blenke, E. O.; Gay, N. J. The Kinetics of Endosomal Disruption Reveal Differences in Lipid Nanoparticle Induced Cellular Toxicity. J. Controlled Release 2025, 386, 114047. 10.1016/j.jconrel.2025.114047.

(51) Son, G.; Song, J.; Park, J. C.; Kim, H. N.; Kim, H. Fusogenic Lipid Nanoparticles for Rapid Delivery of Large Therapeutic Molecules to Exosomes. Nat. Commun. 2025, 16 (1), 4799. 10.1038/s41467-025-59489-5.

(52) El-Mayta, R.; Padilla, M. S.; Billingsley, M. M.; Han, X.; Mitchell, M. J. Testing the In Vitro and In Vivo Efficiency of mRNA-Lipid Nanoparticles Formulated by Microfluidic Mixing. J. Vis. Exp. JoVE 2023, No. 191, e64810. 10.3791/64810.

(53) Protocol: Nuclei isolation from mammalian cells. https://www.takarabio.com/learning-centers/automation-systems/icell8-introduction/sample-preparation-protocols/protocol-nuclei-isolation-from-mammalian-cells (accessed 2026-05-26).

