## Supplementary Information for "Peptide additives reprogram the lipid nanoparticle corona and enhance gene delivery in a serum-free environment for lung epithelium"

**Table S1.** The physical characterization of rLNPs and hpLNPs formulated with pipette mixing

| Sample | Size(nm) | PDI | Encapsulation efficiency (%) |
|---|---|---|---|
| rLNP | 183 | 0.143 | 74 |
| H3 hpLNPs NP 0.6 | 188 | 0.167 | 67 |
| H3 hpLNPs NP 1 | 204 | 0.156 | 35 |
| sH3 hpLNPs NP 0.6 | 198 | 0.129 | 64 |
| sH3 hpLNPs NP 1 | 202 | 0.130 | 23 |

**Table S2.** pKa measurement using TNS assay on various LNP

| Sample | pKa |
|---|---|
| rLNP | 6.62±0.02 |
| sH3 hpLNPs NP 0.9 | 6.63±0.02 |
| sH3 hpLNPs NP 0.9 | 6.64±0.01 |

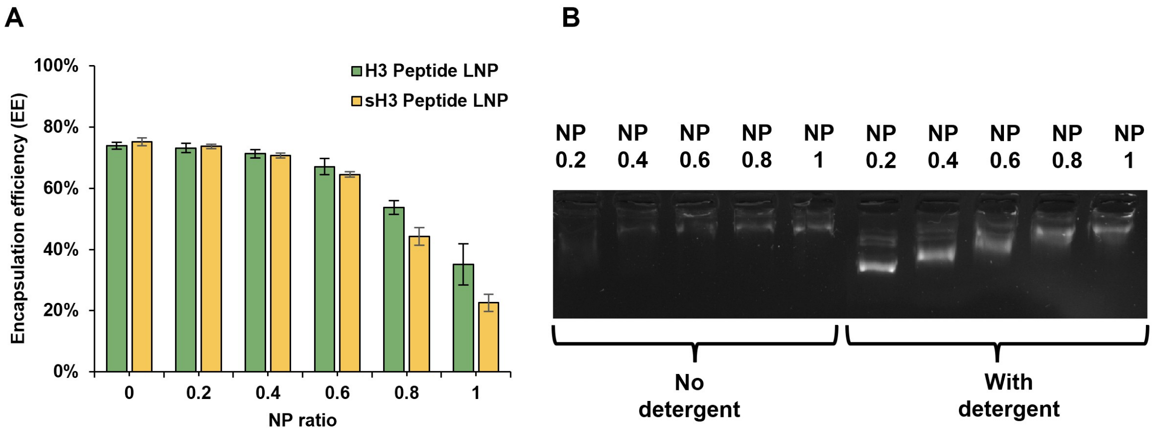


**Figure S1.** Characterization of rLNPs and hpLNPs formed through pipette mixing. (A) Encapsulation efficiency measurements using the PicoGreen assay for hpLNPs with N/P ratios ranging from 0-1. (B) Corresponding gel electrophoresis image of representative hpLNPs with and without detergent disassembly.

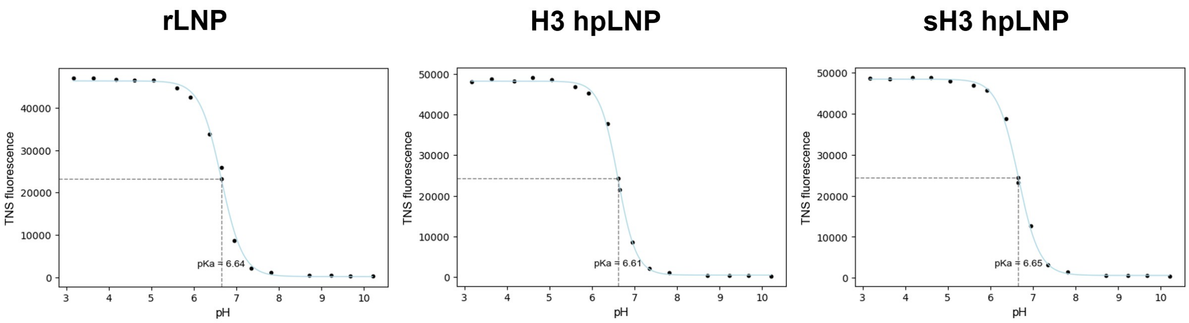


**Figure S2.** Representative pKa measurements using the TNS assay for all types of LNPs.

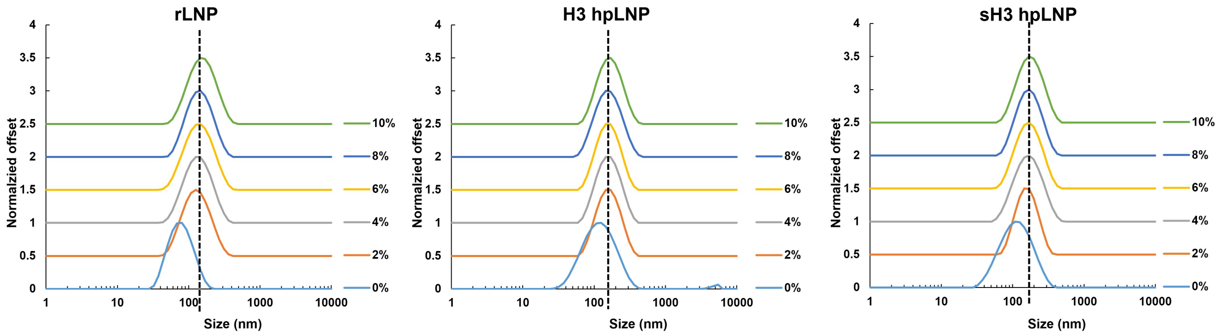


**Figure S3.** DLS size measurements of rLNPs and hpLNPs after 44 h incubation with different quantities of serum-containing DMEM (0-10 %).

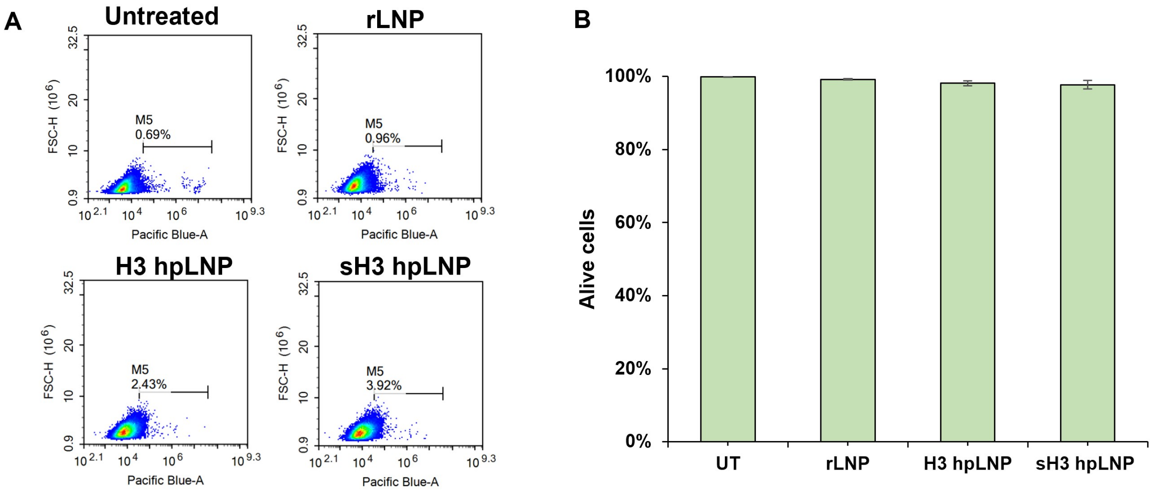


**Figure S4.** Quantification of cell viability using DAPI staining. (A) Representative flow plots of DAPI fluorescence in various LNP dosing condition. The gate in each panel represents DAPI negative cell populations (B) Cell viability of MLE-12 cells 48 h after treatment with LNPs. The data represent the mean ± SD (n = 3).

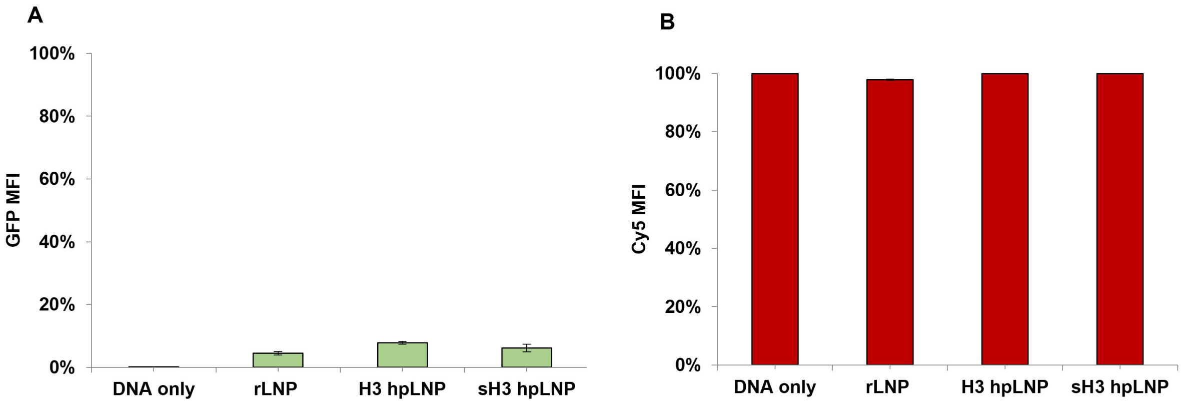


**Figure S5.** *In vitro* MH-S cell transfection and uptake data. (A) GFP expression levels quantified using flow cytometry analysis in MH-S cells treated with various LNPs following preconditioning of LNPs at a serum concentration of 6%. (B) Cy5 signal quantified using flow cytometry in MH-S cells to assess DNA uptake analysis following LNP treatment. Both sets of data represent the mean ± SD (n = 3).

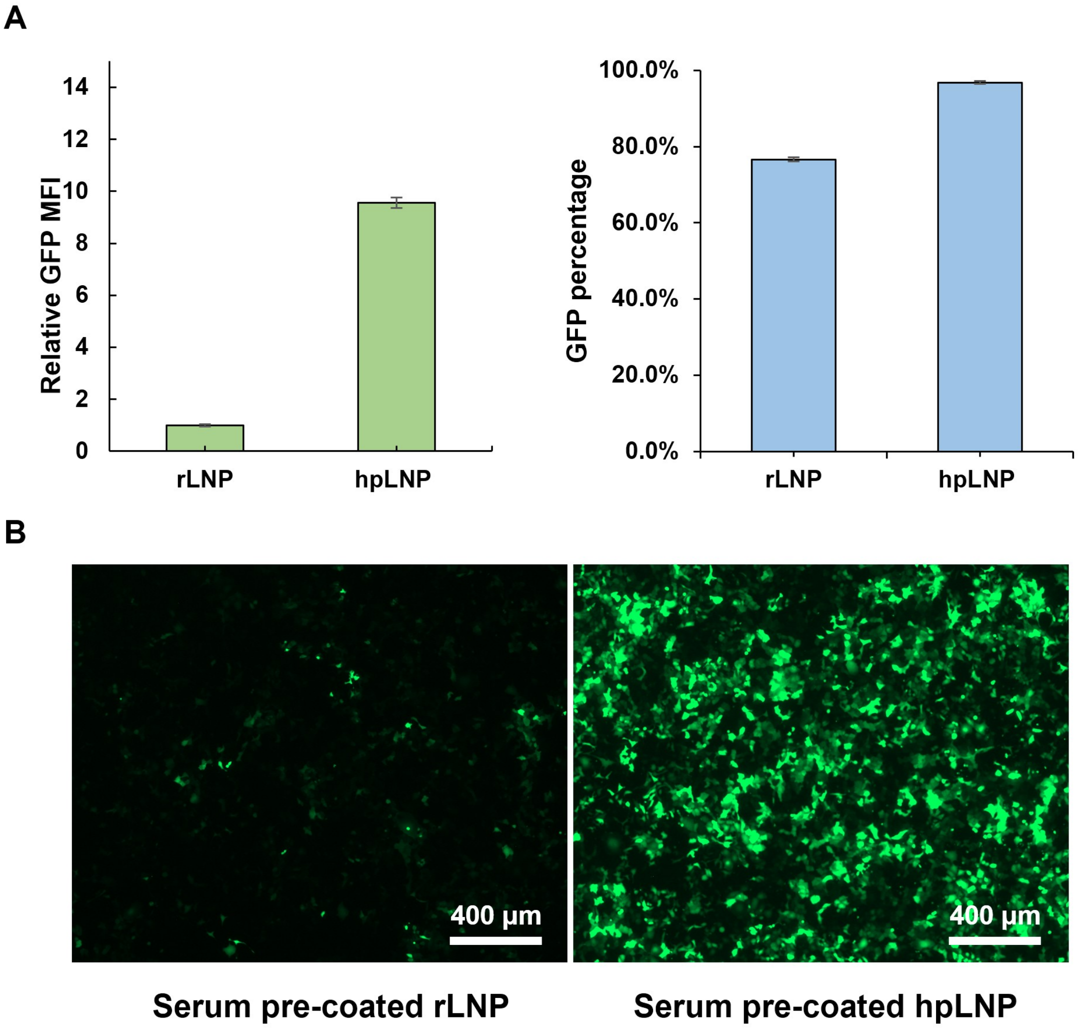


**Figure S6.** *In vitro* MLE-12 cell transfection experiment with serum pre-coated LNPs that were stored for 3 weeks at 4°C prior to transfection. (A) GFP expression levels quantified using flow cytometry. LNPs were stored for 3 weeks at 4°C prior to *in vivo* dosing (n=3). (B) Representative fluorescent images of MLE-12 cells 48 h after transfection with various pre-coated LNPs.

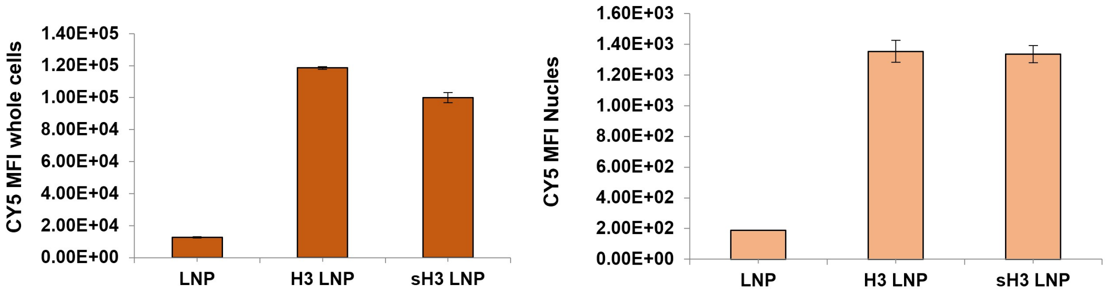


$$Nuclues\ uptake\ efficiency = \frac{Cy5\ MFI_{nuclues} - Cy5\ MFI_{nuclues\ background}}{Cy5\ MFI_{whole\ cells} - Cy5\ MFI_{whole\ cells\ background}} * 100\%$$

**Figure S7.** Nuclear uptake efficiency verification using various stained LNPs. Both cell and nuclear uptake were obtained via flow cytometry (n=3).

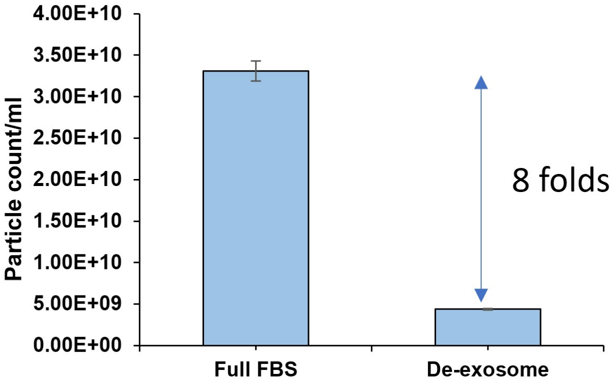


**Figure S8.** Native particle count measurement using a ZetaView NTA device of commercially available exosome depleted FBS.

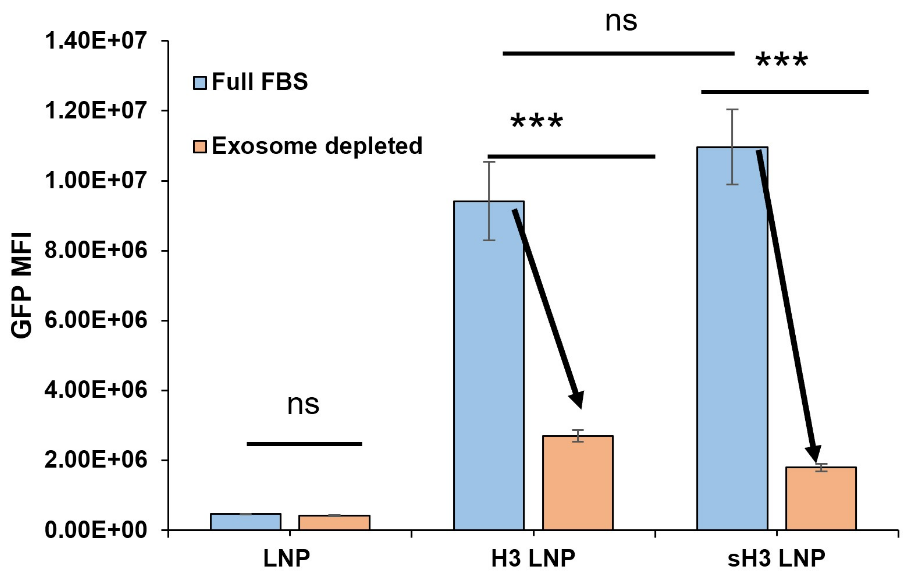


**Figure S9.** Flow cytometry measurement of GFP expression levels following transfection with LNPs that were pre-incubated with normalized protein concentration for 6% serum-containing DMEM media with and without exosomes.

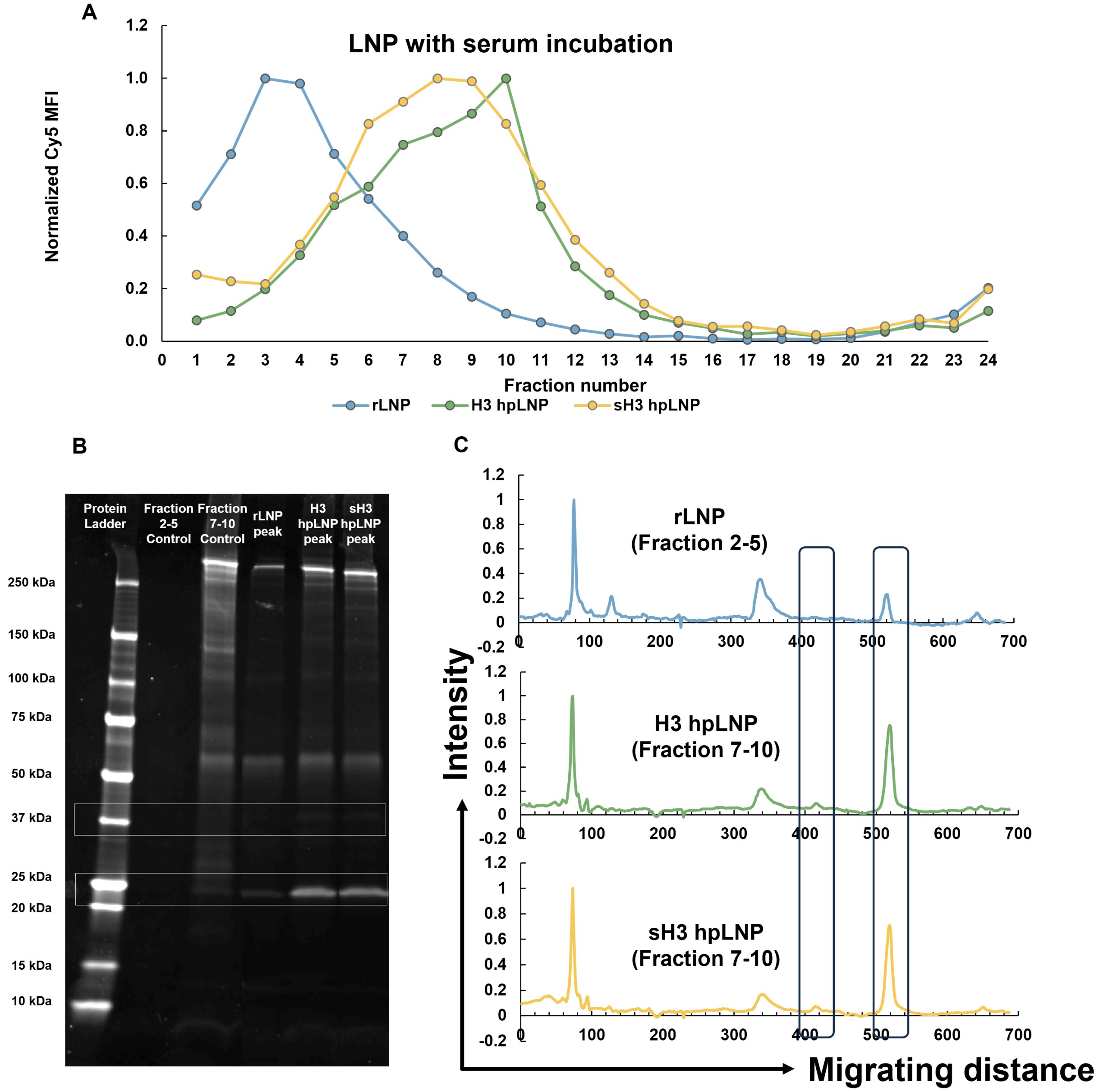


**Figure S10.** Protein corona isolation and analysis of various LNPs. (A) Normalized fluorescent reading of individual fractions containing stained LNPs. Only the 4 fractions containing maximal fluorescence (2-5 for rLNP and 7-10 for hpLNPs) of each LNP were concentrated and isolated for protein gel analysis. (B) SYPR-stained SDS-PAGE. (C) Grayscale analysis of SDS-PAGE data. The background signals were removed using the control fractions in the greyscale analysis and the resulting graph were normalized to the max intensity peak.
